# The impact of package selection and versioning on single-cell RNA-seq analysis

**DOI:** 10.1101/2024.04.04.588111

**Authors:** Joseph M Rich, Lambda Moses, Pétur Helgi Einarsson, Kayla Jackson, Laura Luebbert, A. Sina Booeshaghi, Sindri Antonsson, Delaney K. Sullivan, Nicolas Bray, Páll Melsted, Lior Pachter

## Abstract

Standard single-cell RNA-sequencing analysis (scRNA-seq) workflows consist of converting raw read data into cell-gene count matrices through sequence alignment, followed by analyses including filtering, highly variable gene selection, dimensionality reduction, clustering, and differential expression analysis. Seurat and Scanpy are the most widely-used packages implementing such workflows, and are generally thought to implement individual steps similarly. We investigate in detail the algorithms and methods underlying Seurat and Scanpy and find that there are, in fact, considerable differences in the outputs of Seurat and Scanpy. The extent of differences between the programs is approximately equivalent to the variability that would be introduced in benchmarking scRNA-seq datasets by sequencing less than 5% of the reads or analyzing less than 20% of the cell population. Additionally, distinct versions of Seurat and Scanpy can produce very different results, especially during parts of differential expression analysis. Our analysis highlights the need for users of scRNA-seq to carefully assess the tools on which they rely, and the importance of developers of scientific software to prioritize transparency, consistency, and reproducibility for their tools.

## 1 Introduction

Single-cell RNA-sequencing (scRNA-seq) is a powerful experimental method that provides cellular resolution to gene expression analysis. In tandem with the widespread adoption of scRNA-seq technologies, there has been a proliferation of methods for the analysis of scRNA-seq data^1^. However, despite the large number of tools that have been developed, the majority of analysis of scRNA-seq takes place with one of two analysis platforms: Seurat^2^ or Scanpy^3^. These programs are ostensibly thought to implement the same, or very similar, workflows for analysis^4^. The first step in the computational analysis of scRNA-seq results is converting the raw read data into a cell-gene count matrix *X*, where entry *X*_*ig*_ is the number of RNA transcripts of gene *g* expressed by cell *i*. Typically, cells and genes are filtered to remove poor-quality cells and minimally expressed genes. Then, the data are normalized to control for non-meaningful sources of variability, such as sequencing depth, technical noise, library size, and batch effects. Highly variable genes (HVGs) are then selected from the normalized data to identify potential genes of interest and to reduce the dimensionality of the data. Subsequently, gene expression values are scaled to a mean of zero and variance of one across cells. This scaling is done primarily to be able to apply principal component analysis (PCA) to further reduce dimensionality, and to provide meaningful embeddings that describe sources of variability between cells. The PCA embeddings of the cells are then passed through a k-nearest neighbors (KNN) algorithm in order to describe the relationships of cells to each other based on their gene expression. The KNN graph is used to produce an undirected shared nearest neighbor (SNN) graph for further analysis, and the nearest neighbor graph(s) are passed into a clustering algorithms to group similar cells together. The graph(s) are also used for further non-linear dimensionality reduction with t-distributed stochastic neighbour embedding (t-SNE) or Uniform Approximation and Projection method (UMAP) to graphically depict the structure of these neighborhoods in two dimensions. Finally, cluster-specific marker genes are identified through differential expression (DE) analysis, in which each gene’s expression is compared between each cluster and all other clusters and quantified with a fold-change and p-value.

Seurat, written in the programming language R in 2015, is particularly favored in the bioinformatics community as one of the first platforms for comprehensive scRNA-seq analysis^2^. Scanpy is a Python-based tool that was developed after Seurat in 2017 and now offers a similar set of features and capabilities^3^. Both tools have a wide range of options for analysis and active communities. The choice between Seurat and Scanpy often boils down to the user’s programming preference.

The input to Seurat and Scanpy is a cell-gene count matrix, with two popular packages for count matrix generation being Cell Ranger and kallisto-bustools (kb). Cell Ranger, developed by 10x Genomics, is specifically optimized for processing data from the Chromium platform, providing a solution that includes barcode processing, read alignment (using the STAR aligner^5^), and gene expression analysis^6^. It is popular for its user-friendliness and seamless integration with 10x Genomics data. However, Cell Ranger’s robustness comes with the trade-off of high computational demands, particularly for larger datasets^7^. On the other hand, kb^7;8^ is an open-source alternative to Cell Ranger known for its efficiency and speed^7^. kb-python is a wrapper around kallisto^9^ and bustools^10^, which pseudoaligns reads to produce a barcode, unique molecular identifier (UMI), set (BUS) file, which is then processed into a cell-by-gene count matrix. Utilizing the kallisto pseudoalignment algorithm and the bustools toolkit, kb provides a fast, lightweight solution for quantifying transcript abundances and handling BUS files. This efficiency makes it particularly suitable for environments with constrained computational resources. Additionally, kb is accurate^11^ and stands out for its flexibility, allowing researchers to tailor the analysis pipeline to a broader range of experimental designs and research needs. New versions of each of these packages are periodically released, contributing improvements in algorithm design and efficiency, new capabilities, and integration with new sequencing technologies.

A “standard” scRNA-seq experiment costs thousands of dollars, with exact pricing influenced largely by data size. While it is difficult to provide an exact cost as a result of variability between methods, it is estimated that a typical sequencing kit costs approximated in the range of hundreds to thousands of dollars, and sequencing costs add up to an additional $5 per million reads^12^. The necessary number of reads per cell for high-quality data depends on the context of the experiment, but as an example, Cell Ranger typically recommends 20,000 read pairs per cell for its v3 technologies, and 50,000 read pairs per cell for its v2 technologies^13^. Sample preparation also has substantial costs, often requiring precious patient samples, or maintenance of cell or animal lines for months to years in preparation for experimental analysis. A standard 10x Genomics scRNA-seq experiment sequences tens of millions to billions of reads, with a recommended cell count ranging from 500-10,000+ depending on the context. These estimates do not factor in additional costs including labor, experimental setup, and follow-up analysis. Therefore, it is desirable to try to achieve a middle ground between dataset richness and experimental costs, which requires evaluating the additional information provided by marginal increases in data size.

A typical implicit assumption in bioinformatics data analysis is that the choice among packages and versioning should have little to no impact on the interpretation of results. However, sizeable variability has been observed between packages or versions, even when performing otherwise similar or seemingly identical analyses^14^. The goal of this study is to quantify the variability in the standard scRNA-seq pipeline between packages (i.e., Seurat vs. Scanpy) and between multiple versions of the same package (i.e., Seurat v5 vs. v4, Scanpy v1.9 vs. v1.4, Cell Ranger v7 vs. v6). Additionally, we quantify the variability introduced through a range of read or cell downsampling and compare this to the variability between Seurat and Scanpy.

### 2 Results

## 2.1 Seurat and Scanpy Show Considerable Differences in ScRNAseq Workflow with Defaults

Figure 1 shows the results of comparing Seurat v5.0.2 and Scanpy v1.9.5 with default settings using the PBMC 10k dataset, demonstrating the typical variability to be expected between the two implementations of the “standard” single-cell RNA-seq workflow. Additional pipeline settings run were those with aligned function argument values (Supp Fig 2), identical input data preceding each step (Supp Fig 3), and both aligned function argument values and identical input data preceding each step in the manner of Seurat (Supp Fig 4) and Scanpy (Supp Fig 5).

**Figure 1:**
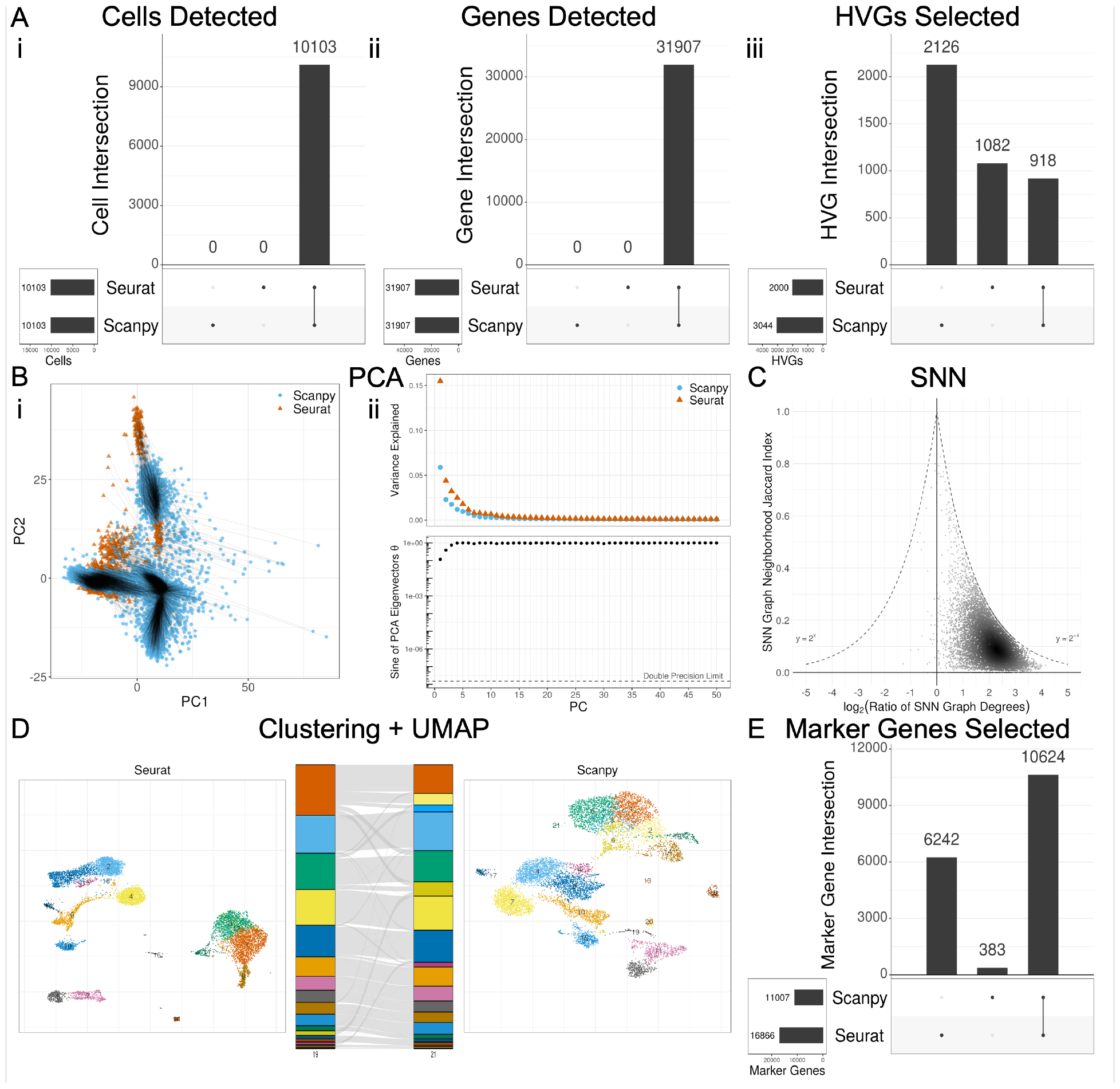
Seurat and Scanpy show considerable differences in scRNA-seq workflow results with default function arguments. (A) Filtering and HVG selection analysis UpSet plots consisting of overlap of sets of cells, genes, and HVGs. (B) PCA analysis through projection onto first 2 PCs, Scree plot comparison of proportion of variance explained, and sine of eigenvectors. Black lines = point mapping between conditions. (C) KNN/SNN analysis through SNN neighborhood Jaccard index and degree ratio (Seurat/Scanpy) per cell. (D) Clustering and UMAP analysis through UMAP plots of each condition, with alluvial plot showing cluster assignment mapping and degree of agreement. Numbers at the bottom of each alluvial plot show the total number of clusters in each group. (E) Differential expression analysis UpSet plot through overlap of all significant (p *<* 0.05) marker genes across all clusters.

There was no difference in cell or gene filtering between the packages after filtering UMIs, minimum genes per cell, minimum cells per gene, and maximum mitochondrial gene content (Fig 1a, i-ii). Furthermore, given the same matrices as input, Seurat and Scanpy handled log normalization identically as well, producing equivalent output (data not shown). However, the programs deviated from their default algorithm for HVG selection, with a Jaccard index (intersection over union between two sets) of 0.22 (Fig 1a, iii). This difference could be resolved either by selecting the “seurat v3” flavor for Scanpy or the “mean.var.plot” algorithm for Seurat (Supp Fig 2a, Supp Fig 5a).

Further differences were observed with PCA analysis, which also yielded different results when run with default parameters. The PCA plots showed noticeable differences in the plotted positions of each cell on the PC1-2 space, although the same general shape of the plot is preserved (Fig 1b, i). The Scree plots also displayed differences, most notably with the proportion of variance explained by the first PC differing by 0.1 (Fig 1b, ii). The eigenvectors demonstrated differences, with the angle between the first PC vectors having a sine of 0.1, the angle between the second PCs having a sine of 0.5, i.e., 30 degrees apart, and PCs 3+ being nearly orthogonal (Fig 1b, ii). All of these changes could be resolved with HVG-set standardization and with the clipping and regression settings prior to PCA adjusted accordingly (Supp Fig 2b, Supp Fig 5b). Seurat, by default, clips values to a maximum of 10 during scaling and does not perform any regression, whereas Scanpy, by default, does not implement clipping and regresses by total counts and percentage of mitochondrial content.

Next, the packages differed substantially in their production of an SNN graph. Both the content and size of each neighborhood per cell differed greatly (Fig 1c). The median Jaccard index between the neighborhood of each cell from Seurat and Scanpy was 0.11, and the median degree ratio (Seurat/Scanpy) magnitude was 2.05. The degree ratio for each was nearly always greater than 1, indicating that Seurat, by default, yields more highly connected SNN graphs than Scanpy. Given that the points in Fig 1c are distributed relatively evenly between 0 and the maximum potential Jaccard index across all degree ratios, it appears that it is not simply the degree difference driving the low median Jaccard index. When aligning the function arguments when generating the SNN graph, there was no qualitative improvement in median degree ratio magnitude, but there was a slight improvement in median Jaccard index (Supp Fig 4c, Supp Fig 5c).

Clustering with default settings also resulted in differences in output, as seen by the discordance in the alluvial plot and the Adjusted Rand Index (ARI) of 0.53 (Fig 1d). Alluvial plots were aligned to maximize cluster alignment and coloring between groups, as to allow visual discordance to correlate with dissimilarity (see Methods). Even when aligning function arguments and input SNN graphs, Seurat and Scanpy demonstrated differences in Louvain clustering (Supp Fig 4d), but were identical in their implementation of the Leiden algorithm (Supp Fig 5d).

UMAP plots visually showed some differences in the shapes of local and neighboring clusters, even when controlling for global shifts or rotations (Fig 1d). For instance, the Seurat UMAP shows that clusters 7 (pink) and 8 (grey) are very distinct from all other clusters on the plot, whereas in the Scanpy UMAP plot, the analogous clusters are located much closer to other clusters, such as 19 (black) and 20 (orange). Comparing neighborhood similarity of KNN graphs constructed from these UMAP data revealed poor neighborhood overlap that modestly improves as the similarity between function arguments and preceding input are aligned (Supp Fig 6). Performing Leiden clustering and subsequent UMAP plotting of these UMAP-derived KNN graphs revealed that the general characteristics of the UMAP plots between packages were maintained, but there were still some considerable irreconcilable differences (Supp Fig 6).

Upon DE analysis, Seurat and Scanpy overlapped with a Jaccard index of 0.62 for their significant marker genes (i.e., the total set of genes with adjusted p-value *<* 0.05 across all clusters), but Seurat had approximately 50% more significant marker genes than Scanpy. (Fig 1e). The difference in significant marker genes is a result of a few differences in default settings between packages. First, each package implements the Wilcoxon function separately, with Seurat requiring tie correction and Scanpy by default omitting tie correction. Additionally, each package adjusts p-values differently by default Seurat with Bonferroni multiple testing correction, and Scanpy with Benjamini-Hochberg multiple testing correction. Finally, Seurat, by default, filters markers by p-value, percentage of cells per group possessing the gene, and log-fold change (logFC) prior to performing the Wilcoxon rank-sum test; Scanpy does not perform this type of filtering without invoking additional functions. Setting the filtering arguments and clusters of Scanpy to be the same as Seurat (filtering, tie-correction, Bonferroni correction) for DE analysis improved the Jaccard index of significant marker gene overlap to 0.73 (Supp Fig 2e), and providing the same cluster assignments further improved the Jaccard index to 0.99 (Supp Fig 4e, i). The remaining 1% of genes differ as a result of differences in logFC calculation discussed later. Setting the methods to be like Scanpy (no filtering, Benjamini-Hochberg) worsened the Jaccard index to 0.38, as a result of the inability to turn off tie correction in Seurat (Supp Fig 5e, i).

When aligning the cluster assignments between groups, further DE analysis can be performed that compares differences in expression levels per gene per cluster. In addition to comparing the sets of significant marker genes across all clusters, the similarity in markers (i.e., genes per cluster after DE analysis and any potential filtering for differential expression) can be compared. As mentioned previously, Scanpy’s lack of filtering means that Scanpy includes all genes in all clusters, even when that gene is minimally- or non-differentially expressed in that cluster; whereas Seurat, by default, includes only a small percentage of genes per cluster based on logFC, p-value, and number of cells expressing the gene in the reference groups (Supp Fig 3e, ii). Applying analogous thresholding to Scanpy vastly reduces the problem, increasing the Jaccard index from 0.22 to 0.92, but not fulling resolving the discrepancy (Supp 4e, ii). Removing all filtering from Seurat fully removes all differences in marker sets between packages (Supp Fig 5e, ii).

Seurat and Scanpy compute logFC differently as well. Comparing each analogous gene per cluster across packages resulted in a concordance correlation coefficient (CCC) of 0.98 and a PCA fit line with a slope of 1, indicating strong correlation across packages. Briefly, CCC measures the agreement between two variables both in terms of correlation and variance. However, observing the scatterplot of logFC values revealed noticeable differences in a large number of values (Supp Fig 3, iii). Specifically, there were a handful of cases (4,109 out of 135,185 markers) where Scanpy predicted a logFC near *±*30 for a gene in a cluster while Seurat predicted a logFC near 0. The reasons for this are elaborated in the Discussion

Regarding adjusted p-value, there were also differences between Seurat and Scanpy (Supp Fig 3e, iv). With default function arguments, Seurat predicted p-values either less than or similar to Scanpy, but never substantially greater. Most p-values were near the maximum of 1, but there was a wide degree of variability. A considerable number of p-values were far from the y=x line, including those below 1e-50 for Seurat but near 1 for Scanpy. 20% of markers had their p-values flip across the p=0.05 threshold between packages, with it being fairly even flipping in either direction (i.e., significant only in Seurat, or significant only in Scanpy). When function arguments were aligned to be like Seurat, virtually all differences in adjusted p-value disappeared (Supp Fig 4e, iv). However, the differences could not be reconciled with Scanpy-like function arguments, due to the lack of ability to toggle tie correction in Seurat’s/presto’s Wilcoxon rank sum calculation.

The comparison of Seurat and Scanpy reveals that, in some but not all cases, the results between programs can be reconciled. There are three classes of possible alignment between functions: aligned by default, aligned when function arguments are matched, and incompatible for alignment. The classification of each function into these classes is shown in Table 1. Supplemental Tables 1 (Seurat) and 2 (Scanpy) go into detail for each step of analysis on the function name, the default arguments, the arguments needed to match the other package as closely as possible, and the parameters unique to that package.

**Table 1:**
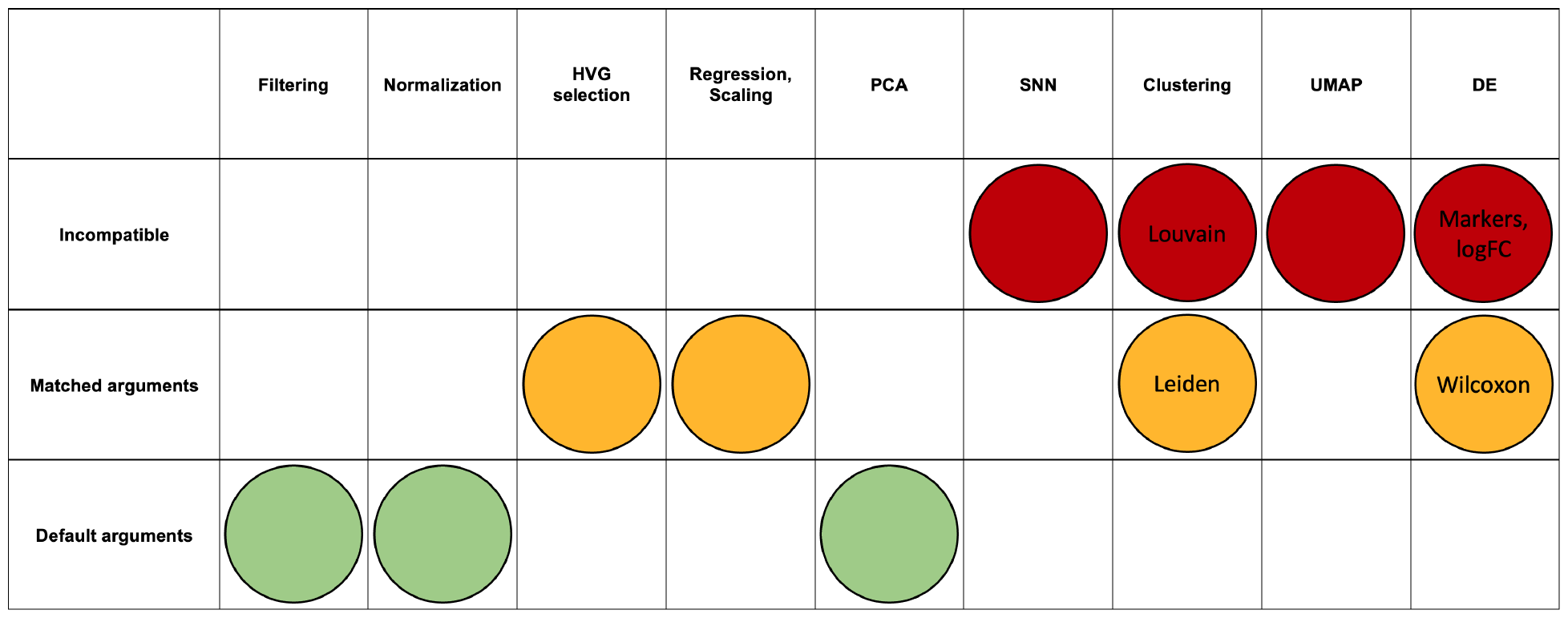
Seurat and Scanpy function agreement for scRNA-seq pipeline. Green = equivalent by default; yellow = equivalent with matched arguments; red = incompatible. HVG = Highly Variable Genes; PCA = Principal Component Analysis; SNN = Shared Nearest Neighbors; UMAP = Uniform Manifold Approximation and Projection; DE = Differential Expression

### 2.2 Read and Cell Downsampling Retain Most Information Compared to Seurat vs. Scanpy Down to Small Fractions of Dataset Size

Given the variability introduced between packages, a natural question that arises is how to benchmark the magnitude of these differences. To this end, we simulated the downsampling of reads and cells before generating the filtered count matrix and compared the differences introduced along a gradient of downsampled fractions to the full-size data. We performed each step of the analysis with default function arguments for the respective package and without aligning input data preceding each step, except for those steps of DE analysis which required input aligning before each step in order to compare marker gene statistics in identical clusters (marker selection, logFC, and adjusted p-value). For each step of analysis, in addition to generating all plots as in Fig 1, we selected a single numeric metric that would capture the degree of variability between groups as follows:

- Cell filtering: Jaccard index of cell sets
- Gene filtering: Jaccard index of gene sets
- HVG selection: Jaccard index of HVG sets
- PCA: Mean of difference in corresponding PC loadings PCs 1-3
- KNN/SNN: Median magnitude of log of SNN degree ratio
- Clustering: ARI
- UMAP: Median Jaccard index of UMAP-derived KNN neighborhoods across all cells
- Marker gene selection: Jaccard index of significant marker gene sets
- Marker selection: Jaccard index of marker sets
- logFC: CCC
- Adjusted p-value: Fraction of adjusted p-values that flipped across the p=0.05 threshold between conditions

The summary for the fraction of downsampling sufficient to achieve results at least as good as the variability between default Seurat vs. Scanpy within a 5% margin is shown in Figure 2. These methods are especially robust to read downsampling, with most steps achieving similar results with less than 5% of the original reads present (Fig 2a). The methods are also robust to cell downsampling, although to a lesser extent, with most methods performing similarly to baseline at less than 25% of the original number of cells (Fig 2b). The step least robust to downsampling for both reads and cells was gene selection; however, given the similarity of HVGs and marker genes to the full-size dataset across downsampled fractions, it appears that the difference in gene sets lies largely in less significant genes. Plots similar to Figure 1 for downsampled fractions of reads and cells, for both Seurat and Scanpy, that generally achieve similar performance to the variability introduced between default Seurat vs. Scanpy can be found in Supplemental Figures 7-10. Metric calculation across downsampled fractions can be found in Supplemental Figures 11-12.

**Figure 2:**
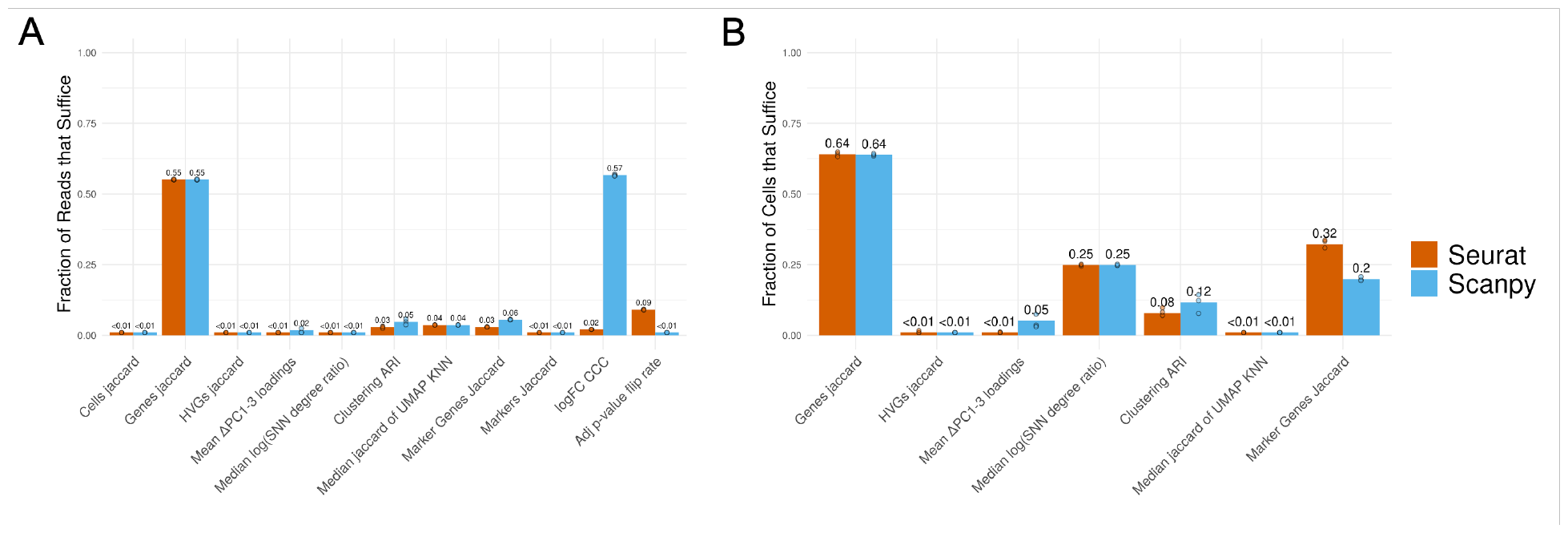
(A) Read and (B) cell downsampling retain most information compared to Seurat vs. Scanpy down to small fractions of dataset size. A minimum downsampled fraction of 0.01 was used as a lower bound. Orange = Seurat; blue = Scanpy.

### 2.3 Package Versioning has Significant Implications with DE Analysis

In addition to package selection (i.e., Seurat vs. Scanpy), package version can also play a role in the interpretation of results. Comparing Seurat v5 to v4, there were sizeable differences in sets of significant marker genes, markers, and logFC estimates (Fig 3a-b). The difference in marker selection arises entirely from differences in logFC calculations and the implications for filtering. The difference in logFC calculation results from a change in the application of pseudocount between versions discussed further in the Discussion. Seurat v4 vs. Scanpy demonstrates the same trend as Seurat v4 vs. v5. The calculation for adjusted p-value remained the same (Fig 3c). Comparing Scanpy v1.9 to the older v1.4 also revealed large differences in sets of significant marker genes and markers as a result of the removal of filtering of markers between releases (Fig 3d). There was no difference in calculation of logFC or adjusted p-value between these versions (Fig 3e-f). And comparing the count matrix generated from Cell Ranger software v7 to Cell Ranger v6 with default settings also revealed differences across all DE metrics (Fig 3g-i). Analysis across Cell Ranger versions showed considerable differences for all steps of the pipeline, as outlined in Supp Fig 13. The primary difference between these commands was the default inclusion of intron counts in the gene count matrix in v7, as opposed to the default exclusion of intron counts in v6. This distinction has implications for UMI filtering and gene per cell violin plots, with Cell Ranger v6 being slightly more restrictive (Supp Fig 14).

**Figure 3:**
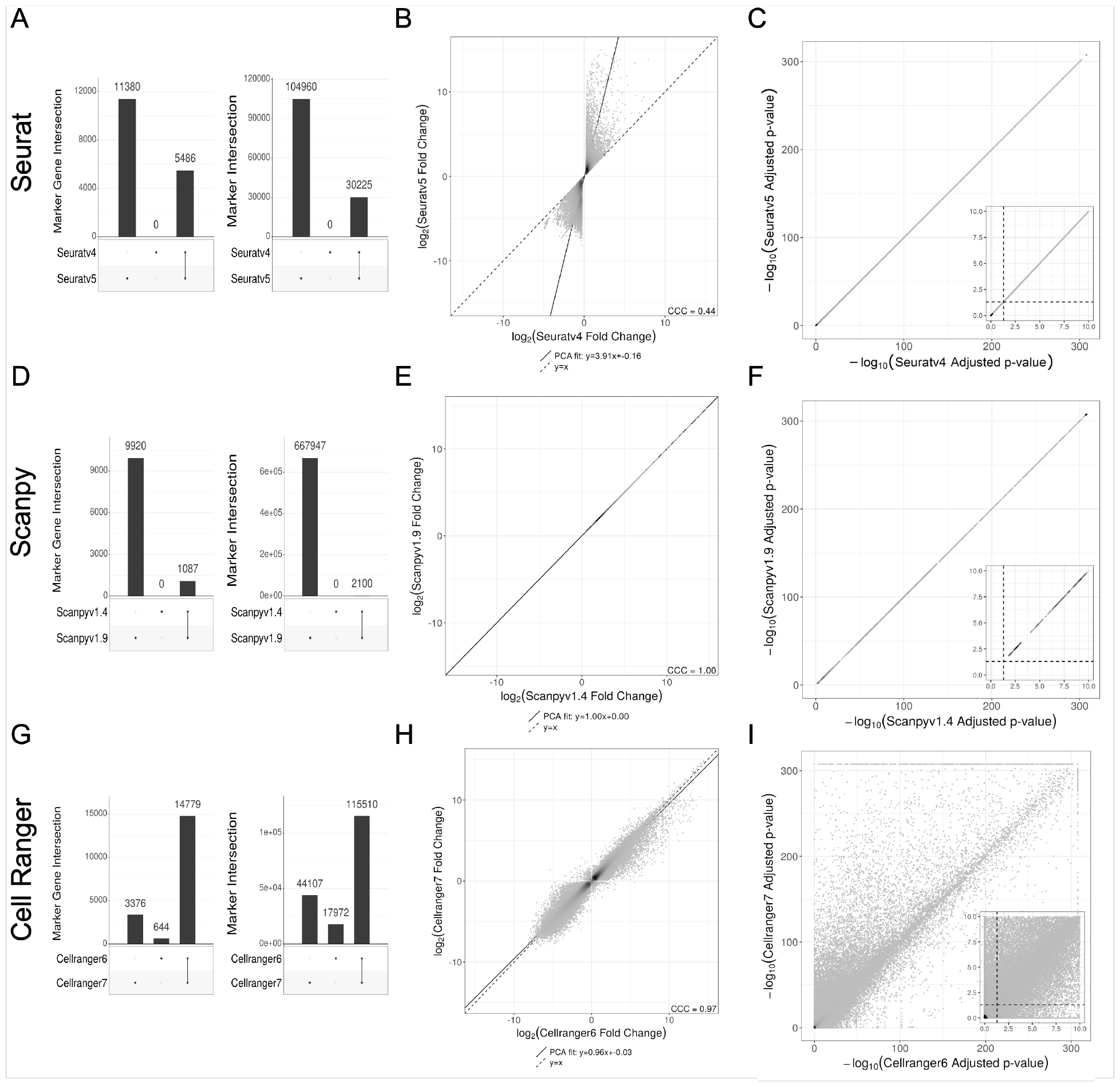
Package versioning has significant implications on DE analysis. (A-C) Seurat v5 vs. v4 differences in (A) significant marker and marker selection, (B) logFC, and (C) Adjusted p-value calculation. (D-F) Scanpy v1.9 vs. v1.4 differences in (D) significant marker and marker selection, (E) logFC, and (F) Adjusted p-value calculation. (G-I) Cell Ranger v7 vs. v6 differences in (G) significant marker and marker selection, (H) logFC, and (I) Adjusted p-value calculation.

### 2.4 Random seeds

The steps of the workflow which involve the influence of randomization are approximate KNN search, graphical clustering with Louvain/Leiden, and UMAP. In order to benchmark the magnitude of difference across packages or data sizes, we ran each of these steps with identical input data and package selection, only varying the random seed applied. The results of comparing each step of Seurat vs. Scanpy with aligned function arguments and identical input preceding each step (Supp figs 4-5) were compared to the variability introduced by differences in random seeds alone (Supp fig 15).

The variations in median Jaccard index and magnitude of log degree ratio of SNN neighborhoods between the same algorithm across random seeds, (0.85 and 0.05 for Annoy, and 1 and 0 for umap-learn/PyNNDescent, respectively), were much less significant than the observed values of 0.27 Jaccard index and 1.61 magnitude log degree ratio observed between and Seurat and Scanpy given identical PCA inputs (Supp Fig 4c, Supp Fig 15a, Supp Fig 15e). The ARI after Louvain clustering between random seeds of 0.96 was significantly higher than that between Louvain implementations in Seurat and Scanpy of 0.85 (Supp Fig 4d, Supp Fig 15b, Supp Fig 15f). The median Jaccard indices of the UMAP-derived KNN across random UMAP seeds, being 0.41 for Seurat and 0.47 for Scanpy, were significantly higher than that when comparing UMAP plots between Seurat and Scanpy with identical input of 0.21 (Supp Fig 6d, Supp Fig 15c, Supp Fig 15g). However, the ARI after Leiden clustering of the same data across random UMAP seeds of 0.64, both for Seurat and Scanpy, was similar to the observed ARI calculated from Seurat vs. Scanpy, given identical PCA and SNN inputs for UMAP of 0.69 (Supp Fig 6d, Supp Fig 15d, Supp Fig 15h). This indicates that despite the higher degree of similarity of UMAP plots generated across random seeds within Seurat or Scanpy, versus UMAP plots generated between the packages, the Leiden algorithm cannot fully capture this similarity.

## 3 Discussion

### 3.1 Matrix generation

There are some assumptions implicit in the discussions above that deserve further scrutiny. First, the assumption that the expression estimates the log-normalized *Y*_*ig*_ derived from the counts *X*_*ig*_ represent accurate measures of expression is frequently taken for granted but is not self-evident. In practice, there is no consensus on how the counts *X*_*ig*_ should be obtained^15;16^. In particular, the pre-processing of single-cell RNA-seq data requires making choices about whether to include in *X*_*ig*_ counts of molecules that are from nascent transcripts, or of molecules that are ambiguous as to their origin from mature or nascent transcripts. These issues are particularly vexing when working with single-nuclear RNA-seq^17^. One approach to “integrating” counts of nascent and mature molecules is to use them together to parameterize models of transcription^18^, raising the question of whether comparisons of parameter estimates in such models are more suitable for assessing differences in transcription between cell types rather than log-fold change estimates based on the *Y*_*ig*_. Moreover, even if one accepts that the (scaled) *Y*_*ig*_ are relevant for measuring expression differences between cell types, it may be that the instability of log-fold change with low expression estimates, which are the norm in single-cell RNA-seq experiments due to the sparsity of data, make them inappropriate as proxies for effect sizes^19^. Finally, implicit in all of the above is the emphasis on effect size, frequently in addition to but sometimes in lieu of the use of pvalues to assess statistical significance. The importance of effect size thresholding in biology research may reflect skepticism of p-values, but may also be popular due to the lack of standards for thresholding allowing for the tuning of thresholds to achieve desired results. Sometimes different thresholds are used within a single paper^20^.

### 3.2 A walk through version histories

Seurat has gone through a series of updates since its initial release in 2015. Seurat v1 introduced tools for the analysis of single-cell RNA-seq data, focusing on spatial localization, clustering, visualization, and identification of markers for cell populations. It was based on the Seurat object, which is a way to store and manipulate single-cell data^2^. Seurat v2 introduced the ability to perform analysis on cell subpopulations across multiple datasets, including the introduction of canonical correlation analysis (CCA)^21^. Seurat v3-v5 further extended the package with a focus on multimodal data integration combining gene expression, spatial transcriptomics, assay for transposase-accessible chromatin with sequencing (ATAC-seq), and immunophenotyping. Seurat v3 helped achieve this with the concept of utilizing anchor cells (cells with a similar biological state across datasets) to merge modalities^22^. Seurat v4 introduced weighted nearest neighbor analysis, an unsupervised approach that determines the weight of each modality in terms of its importance on downstream analysis^23^. Seurat v5 introduced bridge integration inspired by dictionary learning, in which the information of each cell is represented as a linear combination of “atoms” in a dictionary representing information across modalities^24^.

Each major version release was accompanied by additional new features as well. For instance, Seurat v5 includes support for additional assays and datatypes, functionality to explore datasets with large cell numbers without needing to fully load the data into memory, and pseudobulk analysis (aggregating cells within a subpopulation to reduce noise). On a smaller scale, each version of Seurat introduced new function parameters, changes in default settings, and bug fixes. For instance, in the FindAllMarkers function, the default fraction of cells per gene was changed from 0.1 in Seurat v4 to 0.01 in Seurat v5; the default minimum logFC was changed from 0.25 in Seurat v4 to 0.1 in Seurat v5; and the log base was changed from *e* in Seurat v3 to 2 in Seurat v4. While seemingly small, some of these changes had major effects on results, highlighting the importance for users to consider version release statements, and to implement careful environment control for reproducible data analysis.

Scanpy has also had a series of updates since its release in 2017. Throughout development, it references workflows and properties of Seurat. Version 1.0 brought the first major updates, increasing speed and memory efficiency, introducing the Neighbors class, changing PCA implementation through the upgrade of scikit-learn, modifying the input for graph-based tools such as Louvain, and introducing UMAP. Version 1.3 introduced Leiden clustering, batch correction, and calculation of quality control metrics. It also updated the highly variable genes function, and changed behavior to expect log normalized data and not to automatically subset the data structure by HVGs. Version 1.4 introduced filter rank genes groups as a way to provide some filtering functionality after marker gene expression, updated UMAP code, introduced batching in HVG selection, changed solver in the PCA function from auto to arpack, and changed marker gene selection’s t-test implementation to scipy’s. Version 1.5 brought a lot of new changes as a result of AnnData updates, including spatial support, updates to the storage of neighbor and UMAP objects, expanding of scaling and log normalization, PCA implicit centering for sparse matrices (which slightly changes PCA output), and bug fixes to the regression function. Version 1.6 introduced an overhauled tutorial and updated the internals for rank genes groups, including tie correction for Wilcoxon in this function. Versions 1.7 and 1.8 increased spatial support, increased functionality of differential expression analysis, and corrected the output of highly variable gene calculation with seurat v3 flavor and batch key. Version 1.9 introduced usage of Pearson Residuals, fixed multiple bugs with the HVG function (including edge cases with cellranger flavor and the modification of the used layer with seurat flavor), and changed package compatibility with igraph, matplotlib, and scikit-learn versions, among others.

The kallisto quantification tool has also been continually updated since its release in 2015. These changes include the production of BAM and BUS files, implementation of a D-list feature^11^, strand-awareness, workflows to include introns or other custom specifications, the addition of new technologies, and improvement in the pseudoalignment algorithm. kb-python has also gone through a number of changes since its release in 2019, largely reflecting the changes implemented in kallisto and bustools. New versions were accompanied by updated versions in kallisto and bustools, with kallisto v46 with kb-python v0.24.4, kallisto v48 with kb-python v0.27.0, and kallisto v50 with kb-python 0.28.0.

Cell Ranger software was first released in 2016 and has also been continually updated. Early additions to cellranger count included the ability to specify particular genes for analysis or have the ability to exclude some genes for analysis, as well as allowing hard trimming of input FASTQs. Version 3 introduced the EmptyDrops cell calling algorithm called more low RNA content cells, as well as major changes to the VDJ algorithm. Version 4 brought the functionality to trim template switch oligo (TSO) and poly-A sequences from reads, improving alignment and mapping rates. This version also updated the reference transcriptome to 2020-A. Version 5 introduced the ability to include introns in analysis, with the default being false, as well as the multi pipeline to combine gene expression, feature barcode, and V(D)J libraries from a single GEM well. Version 6 introduced support for cell multiplexing, low throughput analysis, and 3’ and 5’ High Throughput kits. It also added the feature that unfiltered feature-barcode matrix files only contain barcodes with at least one read rather than all possible barcodes from the whitelist, shifting the UMI count distribution. Version 7 introduced fixed RNA profiling, modified the batch effect score calculation to normalize and scale with the number of cells in the dataset, and allowed support for FASTQs with quality scores up to the higher upper range of 93 instead of the typical 41. Version 7 also changed the default to include introns to true and removed the necessity to specify the expected number of cells (previously with a default input of 3,000), instead having the ability to calculate a prediction internally by default. The full list of changes introduced with each release can be found on the Cell Ranger website^25^.

### 3.3 Challenges with Seurat and Scanpy

Working across multiple conditions and versions of both Seurat and Scanpy presented programmatic challenges. With Seurat, the FindClusters function with the Leiden algorithm crashed the Rstudio session in the docker container used to run the program as a result of the high-memory intensity of the process. Interestingly, Scanpy implemented Leiden with the same underlying leiden package and produced the same results without the same computational cost. Also, Seurat v4’s FindAllMarkers function was very slow as a result of the slow implementation of the Wilcoxon rank sum test, which was addressed with Seurat v5 with the recommended download of the immunogenomics/presto package. The adjusted p-value calculation outputs are identical between Seurat v5 and v4. Seurat v5 was also not fully compatible with Seurat v4, with some users reporting scripts breaking between versions as a result of changes in assay structure of the Seurat object. Some solutions created by users can be implemented to remedy these discrepancies, but none of these are officially implemented in the package.

Also, running Scanpy 1.4.6 presented issues with package dependency conflicts and a lack of full backward compatibility. Many dependencies did not have upper limits on compatible versions, so these had to be uncovered manually. In our environment, this amounted to matplotlib 3.6.3 and pandas 1.5.3. Additionally, the KNN/SNN method seemingly only worked with umap-learn version 0.5.0 or greater, but the UMAP method seemingly only worked with umap-learn version 0.4.6 or less (which itself required numpy 1.23.0, numba 0.49.1, llvmdev 8.0.0, and pynndescent 0.4.7). Other changes included issues with the calculate qc metrics method, changes to the names of the gene observation column (“n genes” instead of “n genes by counts”) and count observation column (“n counts” instead of “total counts”), use of base e by default during log normalization (instead of base 2) and the need to manually store this base in anndata.uns for the rank genes groups method, storing of KNN/SNN results in anndata.uns (instead of anndata.obsp), and the lack of the “pts” parameter in the rank genes groups method.

### 3.4 HVG selection

The difference in HVG calculation comes entirely from a choice in algorithm, for which the default of each package has an equivalent implementation in the other package. Seurat’s default HVG algorithm is “vst” (equivalent to Scanpy’s “seurat v3” flavor), while Scanpy’s default HVG algorithm is “seurat” (equivalent to Seurat’s “mean.var.plot”). Each of these algorithms calculates the mean and variance of expression values across all cells for each gene, takes some measure to control for the mean-variance relationship, and selects the most variable genes from this list. However, the method for controlling the mean-variance relationship and how they compute their metric for variability in expression differ.

mean.var.plot/seurat bins all genes based on ranked mean expression across all cells in order to control for the mean-variance relationship, where the number of bins *B* is user-provided. For each gene, the dispersion 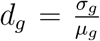 is computed, where *σ*_*g*_ and *µ*_*g*_ are the variance and mean expression of the gene *g* across all cells *i* ∈ *I*, respectively. Within each bin, the mean dispersion 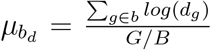 and variance of dispersion 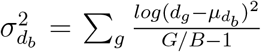 are calculated, where *G* is the total number of genes. A z-score normalized dispersion 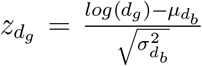 is calculated for each dispersion value based on the mean 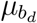 and variance 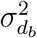 for that gene’s bin. The highly variable genes are either selected based on user-provided thresholds for mean and dispersion values, or by taking the genes with the top *n* dispersion values where *n* is a user-provided fixed cutoff.

vst/seurat v3, rather than binning genes, fits a loess model to the variance and mean expression across all cells of all genes to compute a predicted variance per gene, such that 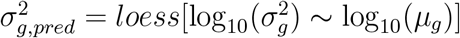. The expression level *X*_*ig*_ for each gene *g* and each cell *i* is “z-score” normalized with its mean expression across all cells and its predicted variance across all cells, such that 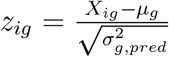. Rather than ranking z-score normalized dispersion values as before, this method then computes the variance of each z-score normalized gene expression values across all cells 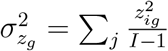 and selects the genes with the top *n* normalized variance values where *n* is a user-provided fixed cutoff (there is no thresholding option as with vst/seurat v3).

### 3.5 PCA

The three methods we used for evaluating PCA similarity projection of cells onto first 2 PCs, variance explained by eigenvalues, and sine of of the angle between eigenvectors all generally correlate with each other. The benchmark displayed on the eigenvector graph is the square root of the double precision limit, which represents a number close to the numerical precision of computers. This benchmark provides a marker to assess when two eigenvectors become so similar as to be technically indistinguishable, but this may be smaller than the smallest difference that is biologically relevant. In all cases where the PCA embeddings were not identical, the sine of even the most similar eigenvectors would never drop below 0.01, even when the PCA plots in 2 dimensions looked reasonably similar.

### 3.6 KNN/SNN

Both Seurat and Scanpy generate two graphs, a KNN and SNN graph, from their nearest neighbors functions. While the KNN graph is a directed graph describing the k nearest neighbors for each cell in PCA space (with a fixed k), the SNN graph is an undirected graph and more broadly groups cells together based on the similarity of their neighborhoods (Supp Fig 16). Unlike the KNN graph, the SNN graph is reciprocal, meaning that if cell A is a neighbor in the SNN graph of cell B, then cell B is a neighbor in the SNN graph of cell A. Additionally, the neighborhood degree (size) is not fixed in the SNN graph, with a wide range of degrees that includes upper bound that can exceed many times the k parameter. When constructing a KNN graph from an SNN graph, edges are connected based on their similarity in KNN space, although how this similarity is defined differs between approaches (Supp Fig 16, 17). Regarding change in neighborhood degree from the KNN to SNN graph, the trend is that hub nodes, where a hub node a node that is a k-nearest neighbor of many other nodes), tend to have a relatively large number of edges due to their involvement in many other neighborhoods. And peripheral nodes, where a peripheral node is a node that is a k-nearest neighbor of few other nodes), tend to have a relatively small number of edges.

Both Seurat and Scanpy use an SNN graph for downstream clustering, and a KNN graph generated from this step is not used in the rest of the standard workflow followed in this study (i.e., clustering, UMAP). When it comes to UMAP, however, the two packages differ. Seurat’s default UMAP function with uwot uses a newly-generated KNN graph from the PCA embeddings built with the approximate KNN graph program Annoy internal to this function for UMAP. On the other hand, Scanpy’s default UMAP function with umap-learn uses the previously-generated SNN graph to generate the UMAP plot (Supp Fig 17).

Seurat defaults to using the Annoy (Approximate Nearest Neighbors Oh Yeah) algorithm from the RcppAnnoy package for computing an approximate solution to the KNN problem^26^. Annoy optimizes the neighbor search space through the construction of multiple binary trees. Each tree recursively divides the dataset with randomly-selected hyperplanes in highdimensional space until the subsets at the leaf nodes contain approximately 100 data points. This approach effectively reduces the number of candidate neighbors to those within the same and neighboring leaves. For each data point, Annoy traverses the trees, collecting data points within the corresponding leaf and some neighboring leaves, calculates distances to all these points, and selects the k closest points as neighbors. This process is repeated for each data point to construct the KNN graph. An alternative algorithm available through Seurat is RANN (R Approximate Nearest Neighbors), which wraps around the C++ library developed by Arya and Mount^27^.

To compute the SNN graph from the KNN graph, Seurat calculates the Jaccard index of neighborhood overlap for each pair of cells and prunes edges that fall below the user-defined prune threshold (default 1/15), with relevant code on the Seurat Github repository^28^. This means that it is not directly relevant whether any two nodes *i* and *j* are in each other’s KNN graphs, but rather whether they share some number of neighbors, above a threshold, overall.

Scanpy by default uses the umap-learn package for handling nearest neighbors search, which itself makes use an exact KNN algorithm for small datasets (either less than 4096 cells, or less than 8192 cells if the distance metric is Euclidean), or an approximate nearest neighbor search using of the NN Descent algorithm through the PyNNDescent package for large datasets to implement KNN graph construction^29^. In contrast to the binary tree-based approach of annoy, NN Descent uses a graph search approach to build the KNN graph. NN Descent initializes a KNN graph with a random projection forest similar to Annoy, and then optimizes this initial guess with graph search. For each node, it randomly selects a candidate node as a potential nearest neighbor, expands the search to all nodes connected to the candidate node with an edge, adds all of these new nodes as new potential candidate notes, and then keeps the k best new nodes as new candidates for further search based on evaluation with a distance metric. It repeats this process for the best candidate node expand the search from the best untried candidate node to all connected nodes, add all expanded nodes as potential candidates, rank by distance to point of interest, and select the top k nodes until there are no untried candidate nodes remaining or a stopping criterion is reached based on minimal improvements. The remaining k nodes (including the self node) are selected as the k nearest neighbor for that node.

To construct the SNN graph from the KNN graph in Scanpy, it makes a connection between each cell *i* and *j* in the KNN graph if and only if either *i* is a k-nearest neighbor of *j* or *j* is a k-nearest neighbor of *i* (excluding the self node), with relevant code on the UMAP-learn GitHub repository^30^. In other words, the SNN graph for Scanpy is the undirected KNN graph. This means that the degree of each node for the SNN graph is bounded below by *k* −1, but is theoretically bounded above by *N* − 1 (where *N* is the number of nodes). This would be the case when a cell is a k-nearest neighbor of all other cells. While this upper bound is essentially never realized with scRNA-seq data, these hub nodes can exist, with degrees in the hundreds for the SNN graph.

For large datasets, both Seurat and Scanpy utilize approximate KNN search algorithms. For each node, an exact KNN algorithm requires searching through each of the other *n* nodes and ranking the distance between points in *d* dimensional space, giving rise to *O*(*d ·n*^2^) time complexity. Approximate KNN algorithms, making use of the fact that a node is less likely to be neighbors with other far-away nodes, can reduce the time complexity to *O*(*d · n* log(*n*)) with techniques such as greedy approaches or binary search trees.

### 3.7 Clustering

Seurat and Scanpy yield different clustering results with the Louvain algorithm, despite following the approach described originally by Blondel et al.^31^. In short, the Louvain method starts by considering each node as an individual community and iteratively reassigns nodes to maximize modularity, thereby clustering nodes based on the density of intra-community edges. This process of local optimization is followed by grouping the formed communities into a new, reduced network, where each community becomes a node, and the steps are repeated. The algorithm continues until it achieves maximal modularity, effectively revealing the hierarchical community structure within the network at a given resolution. The size of the resulting network can be tuned with the resolution parameter. Interestingly, the packages approach more similar results using their differing default resolution arguments (1.0 for Seurat, 0.8 for Scanpy) rather than the same resolution parameter (Supp Figs 3d, 4d),

Seurat has developed its own implementation of the Louvain algorithm, encapsulated within the ‘RunModularityClusteringCpp’ function. On the other hand, Scanpy leverages the ‘vtraag/louvain-igraph’ package. The difference in outcome results from the differential implementations of the optimizers in C++ that perform the local moves to maximize modularity. In particular, during the moving step of the algorithm, Seurat’s implementation considers only local communities for possible moves, while Scanpy considers all communities. Scanpy’s implementation also provides the possibility to move nodes to empty communities as an option to escape local minima. Each function utilizes different pseudo-random numbers for the influence of random seeds, such as in the order of nodes considered for each moving step, although the influence of randomness in each algorithm is small (i.e., affects ARI by *<* 0.02) (Supp fig 15). Both approaches involve iterating the algorithm until zero nodes move, suggesting an optimal arrangement has been reached, but Seurat additionally implements a user-provided upper limit on number of iterations. And another way that the packages differ is in their calculation of the quality score. Both quality scores are based on modularity. However, Seurat’s implementation compares a cluster’s score to what would be randomly expected with *quality* = *quality* –*resolution* * (*total edges community*)^2^, and further normalizes the score by the total weights of the whole network to control for cluster size when comparing scores. Scanpy, on the other hand, directly calculates the expected number of intracommunity edge by chance (rather than implying a relationship with the square of total edge weights in that community like Seurat), and thus adjusts the quality score with *quality* = *quality* − *resolution* * *expected edges community random*. Scanpy does not incorporate normalization by total network size.

### 3.8 UMAP

The differences between the underlying packages that implement UMAP for Seurat (uwot) and Scanpy (umap-learn) have been addressed in the uwot documentation^32^. The magnitude of visual differences outlined in the examples discussed on this page are similar to those observed with the PBMC 10k dataset used in this study. The source of these differences includes differences in initialization - while both packages use spectral initialization when only one component is present, they differ in initialization approaches when more than one component is present. In these cases, uwot falls back to PCA, while umap-learn uses meta-embedding of the separate components. Additionally, as discussed earlier, uwot internally constructs a new KNN graph (exact if dataset size *<* 4096 observations, Annoy otherwise also implemented by RcppAnnoy) from the PCA embeddings, whereas Scanpy feeds in the SNN graph generated from the previously-run nearest neighbors method for umap-learn. For umap-learn, if it is not provided a nearest neighbor graph (which would be the case if sc.pp.neighbors is not run), the package by default will generate a KNN graph internally with the package PyNNDescent (without as strong of a need to run PCA beforehand if not done already, with the same threshold of 4096 observations for exact vs. approximate implementation, and with PyNNDescent rather than Annoy for when approximate KNN is used). In summary, the differences in UMAP plots can likely be mostly explained by the different approaches of initialization, different nearest neighbor graphs (KNN with Annoy for uwot, SNN with UMAP by default for umap-learn), and random variation introduced with different random seeds.

Interestingly, when overlaying Louvain/Leiden clusters on the UMAP plots, Seurat tends to display more examples of overlapping cluster regions on UMAP space compared to Scanpy. This behavior is likely due to the use of a separate nearest neighbors graph for clustering vs. UMAP generation by Seurat, whereas Scanpy uses the same nearest neighbors graph for each of these. Within Seurat, using Louvain with multilevel refinement (algorithm 2) can reduce this overlap in Seurat clusters in some cases.

### 3.9 Marker selection

For differential expression testing, each package performs logFC and p-value calculations for each gene, comparing each cluster to all other combined clusters. Seurat internally possesses the ability to perform other p-value corrections besides Bonferroni, such as Benjamini Hochberg as Scanpy uses, but does not provide a function parameter to change the correction method; however, this can be performed directly in R. Only Seurat’s DE function also possesses the ability to internally filter genes by logFC, percentage of cells expressing that gene in either comparison group, and p-value, which it does by default; Scanpy requires a separate function for filtering the results of DE after it has been performed, and no way to filter by p-value, although these operations too can be easily performed directly in Python. Without an additional filtering step, Scanpy does not filter any genes during the marker gene identification process. But while each package’s respective marker identification and adjusted p-value calculation can be made identical, the sets of significant marker genes across all clusters do slightly differ when filtering by logFC as a result of each package’s different implementation of logFC calculation.

### 3.10 logFC

Log-fold change is defined as the logarithm of the ratio of expression values between groups, taking the following form

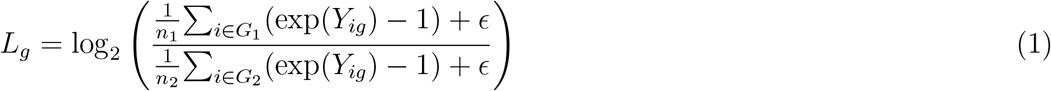

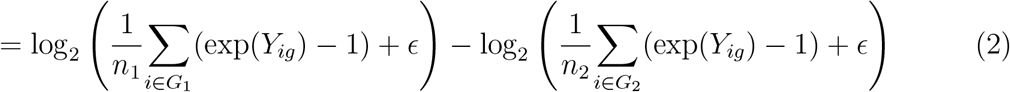

where *Y*_*ig*_ is the log-transformed expression value for cell *i* and gene *g* derived during the log-normalization step of a standard pipeline (i.e., *Y*_*ig*_ = *ln*(*X*_*ig*_ + 1), where *X*_*ig*_ is the raw expression value for cell *i* and gene *g*), *G*_1_ and *G*_2_ are the indices for two groups of cells, *n*_1_ and *n*_2_ are the numbers of cells in the respective groups, and *ϵ* is a very small pseudocount to avoid *log*(0) or *div*(0) errors.

In a recent survey of methods for finding marker genes from single-cell RNA-seq data^33^, the authors urge that “extreme care should be taken when comparing the log fold-changes output by Seurat (version 4) and Scanpy” because the programs are using different formulas for the calculations. Specifically, the Seurat v5 calculation is given by

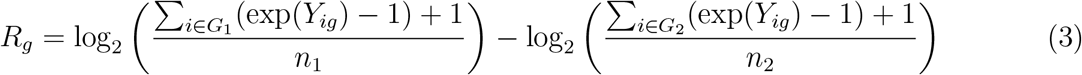

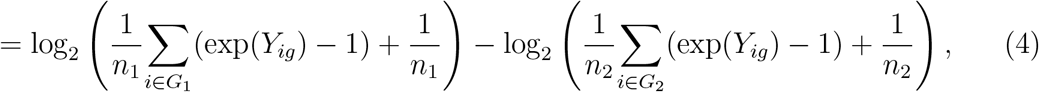

The relevant code can be found on the Seurat GitHub repository^34^.

In contrast, the Scanpy calculation is

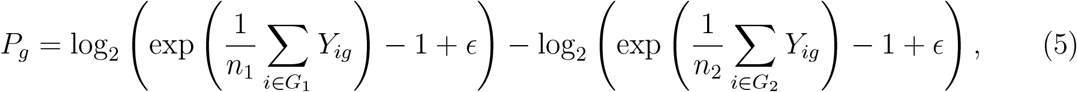

where *ϵ* = 10^−9^. The relevant code can be found on the Scanpy GitHub repository^35^. There are two noteworthy differences in the calculations between packages. The first difference is in the pseudocount, which is 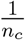 for Seurat (where *c* is the group identity), while it is *ϵ* = 10^−9^ for Scanpy. Seurat incorporates cluster size into its pseudocount, meaning that the pseudocount will increase for smaller clusters (in the range of 0.0001 to 0.1 for a typical dataset). This pseudocount-cluster size dependence means that both the expression level and cluster size will affect the calculated value, such that two clusters with equal mean expression levels for a gene will have a non-zero logFC if the cluster sizes are different. The second difference is that Seurat reverses the log transform *before* calculating the mean across all cells, while Scanpy reverses the log1p transform *after* calculating the mean. In other words, Scanpy erroneously computes exp(*arithmetic mean*(*Y*_*ig*_)) − 1, rather than the correct *arithmetic mean*(exp(*Y*_*ig*_ − 1)).

Seurat v4 calculates the logFC similarly to Seurat v5, but with the crucial difference that the pseudocount is added *after* dividing by total cells in Seurat v4, whereas the pseudocount is added *before* dividing by all cells in Seurat v5, yielding the following equation for logFC in Seurat v4

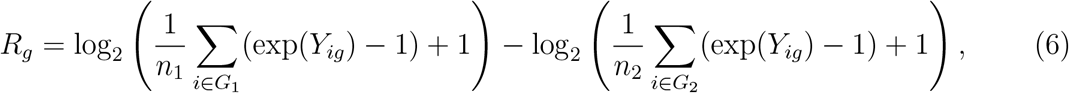

This difference effectively means that the added pseudocount in Seurat v4 is 1, while the added pseudocount in Seurat v5 is 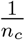. This explains the difference in logFC calculation between Seurat versions. The relevant code can be found on the Seurat GitHub repository under the version 4 release^36^.

It is the difference in handling of the pseudocount ( 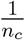 for Seurat v5, 1 for Seurat v4, *ϵ* for Scanpy) which contributes to the large slant when comparing analogous logFC values between Seurat v5 and v4 (Fig 3a), as well as the presence of outliers near (*±*30, 0) when comparing Seurat to Scanpy (Supp Fig 3g). The relatively large pseudocount in Seurat v4 drives all fold-change ratios toward 1, or all log-fold changes toward 0, while Seurat v5’s relatively small pseudocount does not significantly sway its log-fold change values, thus generally trending Seurat’s logFC values to be smaller than Scanpy’s. And between Seurat and Scanpy when a gene is not expressed in one of the groups, rather than treating logFC as *±*∞ as would be technically correct, the pseudocount treats these edge cases differently. As an example, let’s simplify the logFC calculation to 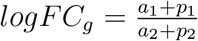, where *a*_1_ and *a*_2_ are the mean unlogged expression values for logFC calculation, and *p*_1_ and *p*_2_ are the pseudocounts, respectively. For Seurat, let’s set *p*_1_ = 0.01 (a reasonable value for a small cluster), and *p*_2_ = 10^−4^. If the expression in group 1 is 0 and the expression in group 2 is 1, then rather than setting the numerator to 0 (and thus providing a logFC of −∞), Seurat would calculate logFC as 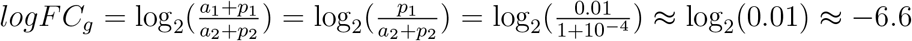. However, Scanpy would make the calculation as 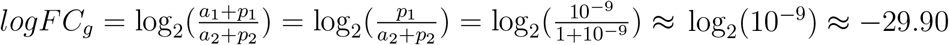 (and similarly *logFC* ≈ +29.90 if groups 1 and 2 are swapped).

Additionally, the values of *Y*_*ig*_ that appear in each equation are not simply *log*1*p*(*X*_*ig*_) as discussed above. Rather *Y*_*ig*_ is typically normalized before being logged, obtained by dividing the raw count *X*_*ig*_ by the total number of counts in cell *i*, and then multiplying by the number 10, 000 (user-defined in Seurat, fixed in Scanpy). Specifically, 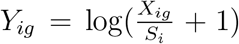 where 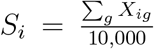 . Thus, 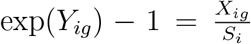, and 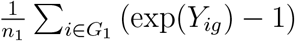 (respectively 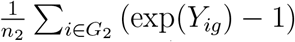 is the arithmetic mean of 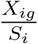 computed across the cells in *G*_1_ (re-spectively *G*_2_).

Seurat computes the log of the arithmetic mean 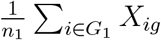 (in what follows we focus on *G*_1_, although the claims and results hold for computations with respect to *G*_2_ as well), which makes sense as it can be understood to be the maximum likelihood estimate (MLE) for the mean of a negative binomial distribution, namely the distribution for the molecule counts *X*_*ig*_. The modeling of the *X*_*ig*_ with a negative binomial distribution represents an assumption about the data in an experiment, but is justifiable^37^. The arithmetic means computed in *R*_*g*_ are not, however of the *X*_*ig*_. As noted above, they are of 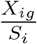, whose distribution will depend on the cell depths *S*_*i*_; moreover the values 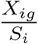 are not integers. Thus, the use of the MLE for the negative binomial distribution may not be appropriate, although in practice the log-fold change is computed for cells in two distinct cell types, and therefore the sets *G*_1_ and *G*_2_ will be homogeneous leading to 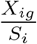 being (approximately) a constant multiple of the *X*_*ig*_. When this is the case, the sample mean 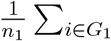 is likely to yield a good estimate of the scaled mean for the negative binomial distribution of the *X*_*ig*_.

However, Scanpy computes the log of 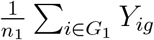, which is equivalent to the log of the geometric mean of exp(*Y*_*ig*_), i.e 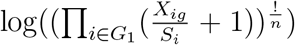. The log of geometric mean is the MLE for the mean of log-normal distributed data, so it makes sense to use the geometric mean if one assumes that the exp(*Y*_*ig*_) are log-normally distributed, however the authors of Scanpy explain that this is not the case, writing^4^ that “scRNA-seq data are not in fact log-normally distributed”. Moreover, there is an arithmetic error in the computation of *P*_*g*_, evident in the subtraction of 1 after computing the geometric mean in the *P*_*g*_ formula 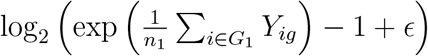. The subtraction is intended to adjust for the fact that the geometric mean is computed for 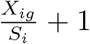 and not 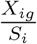; presumably this was a matter of convenience as the *Y*_*ig*_ are stored in the anndata object that Scanpy uses for other purposes. However, while the arithmetic mean is linear, the geometric mean is not, and in general 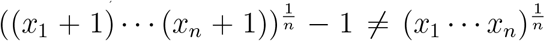. Again, in most cases, the homogeneity of the groups *G*_1_ and *G*_2_ comes to the rescue as 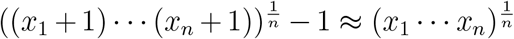 when the *x*_*i*_ are close to each other. The formula *P*_*g*_ also includes a pseudocount to avoid an error in the case when the *Y*_*ig*_ are all equal to zero for a group (and thus the attempted evaluation of the logarithm of zero), but in the case of *P*_*g*_ the pseudocount used is *ϵ* = 10^−9^, thus avoiding the problems with the +1 pseudocount in *R*_*g*_.

Interestingly, Seurat’s logFC calculation is much more stable to reduced read counts than Scanpy’s. Given the general correlation between Seurat’s and Scanpy’s logFC calculations at baseline, it would be more accurate to compute logFC for a dataset with Seurat if hoping to recapitulate results of a larger dataset, even if hoping to mimic the Scanpy logFC calculation.

### 3.11 Adjusted p-value

As the DE output of both Seurat and Scanpy contains p-values and users may use the adjusted p *<* 0.05 cutoff to select genes for further analyses, we compare the p-values to compare the DE results from the standard workflow (Supplemental Figure 9A). Despite using the more conservative Bonferroni correction, Seurat tends to report more significant p-values than Scanpy, which uses the Benjamini-Hochberg correction. The difference in adjusted p-value calculation between packages nearly entirely boils down the required use of tie-correction by Seurat, in contrast to the omission of tie-correction by default in Scanpy. Both Seurat, directly or through immunogenics/presto, and Scanpy perform the Wilcoxon rank sum test in essentially the same way. To briefly review the algorithm, for each gene, it ranks the cells by expression value of that gene, adjusting for ties if indicated. It then loops through the clusters, comparing each cluster to a reference group (by default, all other clusters combined); calculates the mean and standard deviation gene expression for the selected gene in each group across all cells; and sums the ranks of the cells for the selected gene expression values within each group. For the selected cluster, it calculates the standard deviation of all ranks, adjusts the rank sum by the sum expected when no difference is present (dependent on the size of the selected cluster and the size of the reference group) dividing by the standard deviation of ranks, and computes a two-tailed p-value based on this adjusted rank sum score. The use of presto for DE calculations including Wilcoxon rank sum test, new to Seurat v5, reduces the calculation time per run by approximately 80-90%, while providing identical results to the method built into Seurat.

### 3.12 Downsampling

Most results observed with the PBMC 10k dataset are also similarly observed when running the same analysis on the PBMC 5k dataset. One exception, however, is that the number of reads needed to preserve a similar degree of variability introduced by Seurat and Scanpy in the PBMC 5k dataset is nearer 10-25% downsampled, rather than the 1-5% observed with PBMC 10k. Given that the PBMC 10k dataset has approximately five times more reads than the PBMC 5k dataset, it is possible that the fraction of necessary reads to preserve information depends not only on the fraction of total reads, but additionally on the number of reads after downsampling, the number of cells, and other specifications of the dataset. Further datasets with a range of sequencing technologies, cell sample sizes, sequencing depths, and tissue pathologies can be tested to improve the robustness and quantitativeness of these results, although even these two datasets alone provide evidence that at least some datasets are large enough that technical noise between methods can outweigh significant downsampling.

With the downsampling analysis, the point of comparison for the degree of information preservation was always the variability introduced between Seurat and Scanpy. Without extensive biological validation, it is not clear how much the observed differences between the packages, or between versions within a package, matter in practice. However, even without validation, two points are made clear. The first point is that the variability introduced between packages is appreciable for most steps of the pipeline. The second point is that given that similar information is preserved with a small fraction of the original reads, it is true either that the difference in results between default Seurat and Scanpy is significant in the interpretation of biological analysis (in which case the acceptable degree of downsampling is overestimated in this paper), or that similar results can be achieved in some cases with a small fraction of the reads or cells.

## 4 Conclusion

This study lays the foundation for comparing similarity in the outputs of each step in a standard scRNA-seq analysis. While we have illustrated differences between scRNA-seq software tools using 10x Genomics PBMC datasets that are standard for benchmarking, our comparisons can be readily conducted on other datasets. Future efforts could include the broadening of this study to further packages, package versions, and datasets. Additional metrics involved in scRNA-seq can be analyzed, including those in related workflows such as spatial transcriptomics and scATAC-seq. Datasets with various sequencing depths, cell sample sizes, and tissue sources could be studied in order to better understand the necessary dataset sizes needed to capture relevant information across different conditions. Another avenue that was not explored in depth was the effect of argument choice on the divergence of output among conditions. One study found some deviations from default arguments in Seurat to particularly impact clustering results such as number of dimensional reductions to use, k-parameter of KNN, prune parameter of SNN, and resolution parameter of clustering^38^. It would be interesting to apply this analysis to other output metrics, such as UMAP and DE, and see if the impact of these arguments propagates across packages and versions. And applications of this workflow could be applied to biological workflows with well-established ground truths in order to determine to what extent these discrepancies lead to differences in biological interpretation.

We provide guidelines for best practice that can mitigate the differences observed in this study. For developers, maintaining backward compatibility is crucial. Ideally, a unified backend framework can significantly minimize discrepancies between frontend tools. Default function arguments should be justified, especially if they deviate from other popular choices in the field. Announcing all changes to function parameters and default values can provide users to easily understand how analytical steps differ between versions. For users, careful programming environment setup with tools such as virtual environments, Anaconda, Docker, and Google Colab is important for reproducibility. The selection of function arguments should be intentional. A single pipeline should be established within each study to avoid introducing some of the discrepancies discussed in this study. And one should interpret results across studies with caution when different packages, versions, and function arguments are utilized.

In conclusion, we establish a series of methods for assessing the similarity in function of each step of a standard scRNA-seq analysis pipeline from filtering to DE. We highlight how Seurat and Scanpy possess sizeable differences in the way they perform analysis with default settings, which can only be partially reconciled by aligning function arguments. These differences amount to the variability introduced when downsampling reads to less than 5%, or when downsampling cells to less than 20% in the dataset analyzed. Version control of packages involved in count matrix generation and analysis can also have an impact on downstream analysis, especially without careful consideration of changes in behavior across versions. Consistent package selection, thoughtful argument choices, and intentional version control must be practiced in order to achieve accuracy and reproducibility in scRNA-seq analysis.

## Acknowledgements

This work was supported in part by NIH 5UM1HG012077-02. D.K.S. was funded by the UCLA-Caltech Medical Scientist Training Program (NIH NIGMS training grant T32 GM008042). We thank Bernadett Gaál for the feedback of dedicating a study to package comparisons in the scRNA-seq workflow. We thank Tara Chari for providing feedback and assisting with data management. The authors acknowledge the Howard Hughes Medical Institute for funding A.S.B. through the Hanna H. Gray Fellows program. We thank the Caltech Bioinformatics Resource Center for providing computing resources during the development of the project.

## Author Contributions

Work on this paper was led by J.R., who implemented the comparisons, produced the results, and drafted an initial version of this manuscript. The project emerged from discussions among various combinations of the authors: J.R., L.M., P.H.E., K.J., L.L., A.S.B., S.A., D.K.S, N.B, P.M., L.P. The methods for benchmarking and comparisons of results were developed by J.M.R., L.M., L.P.; Software was written primarily by J.M.R. with the help of L.M.; Formal analysis and investigation was conducted by J.M.R., L.M., L.P.; Writing – Original Draft, J.M.R.; Writing – Review & Editing, J.M.R., L.M., P.H.E., K.J., L.L., A.S.B., S.A., D.K.S, N.B, P.M., L.P.; Visualization, J.M.R., L.M., L.P.; Funding Acquisition, L.P.; Resources, L.P.; Supervision, L.P.

## STAR Methods

### KEY RESOURCES TABLE

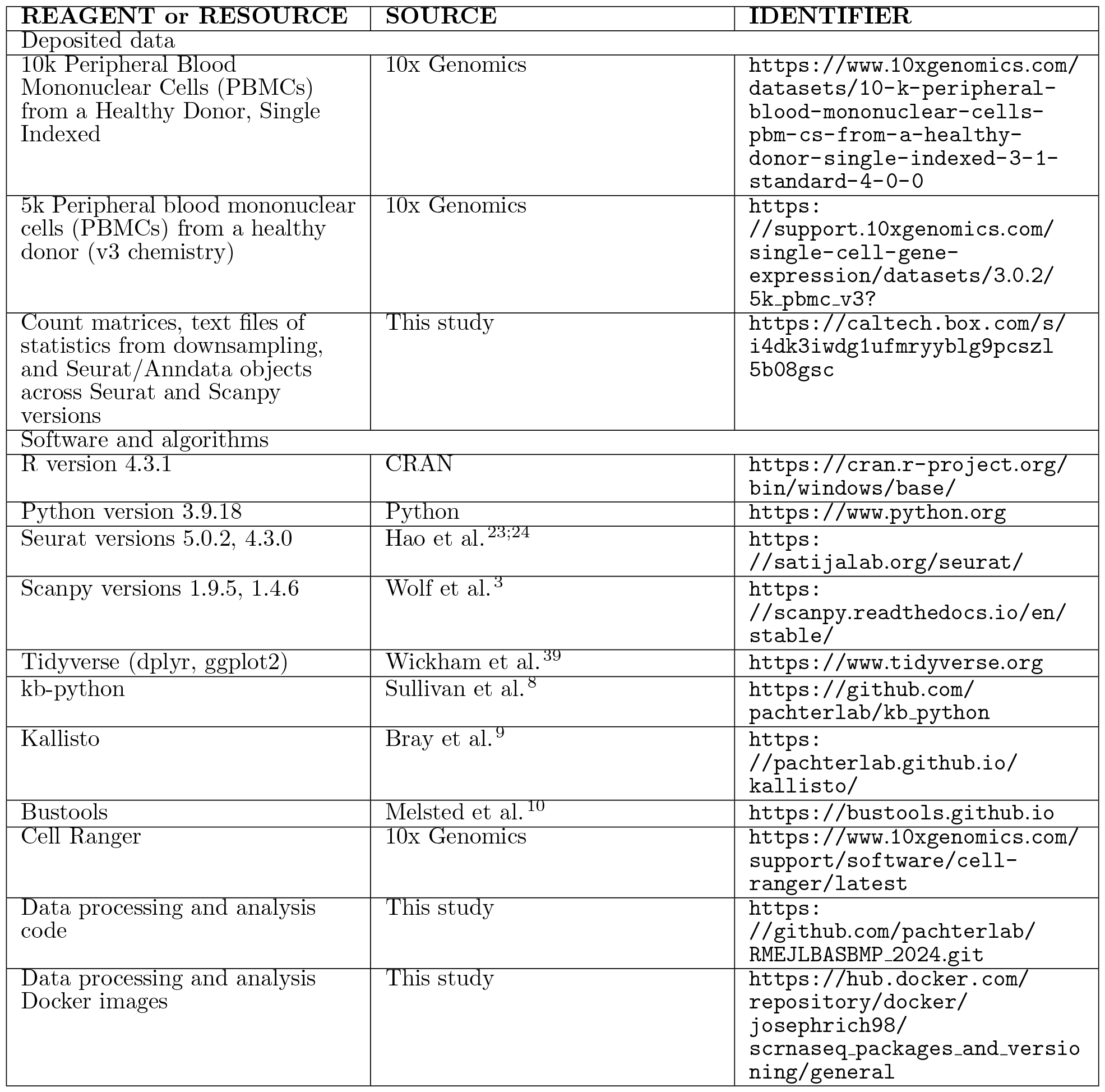

| REAGENT or RESOURCE | SOURCE | IDENTIFIER |
| --- | --- | --- |
| Deposited data |  |  |
| 10k Peripheral Blood Mononuclear Cells (PBMCs) from a Healthy Donor, Single Indexed | 10x Genomics | https://www.10xgenomics.com/datasets/10-k-peripheral-blood-mononuclear-cells-pbm-cs-from-a-healthy-donor-single-indexed-3-1-standard-4-0-0 |
| 5k Peripheral blood mononuclear cells (PBMCs) from a healthy donor (v3 chemistry) | 10x Genomics | https://support.10xgenomics.com/single-cell-gene-expression/datasets/3.0.2/5k_pbmcs_v3? |
| Count matrices, text files of statistics from downsampling, and Seurat/AnnData objects across Seurat and Scanpy versions | This study | https://caltech.box.com/s/i4dk3iwdg1ufmryyblg9pcszl5b08gsc |
| Software and algorithms |  |  |
| R version 4.3.1 | CRAN | https://cran.r-project.org/bin/windows/base/ |
| Python version 3.9.18 | Python | https://www.python.org |
| Seurat versions 5.0.2, 4.3.0 | Hao et al.<sup>23;24</sup> | https://satijalab.org/seurat/ |
| Scanpy versions 1.9.5, 1.4.6 | Wolf et al.<sup>3</sup> | https://scanpy.readthedocs.io/en/stable/ |
| Tidyverse (dplyr, ggplot2) | Wickham et al.<sup>39</sup> | https://www.tidyverse.org |
| kb-python | Sullivan et al.<sup>8</sup> | https://github.com/pachterlab/kb-python |
| Kallisto | Bray et al.<sup>9</sup> | https://pachterlab.github.io/kallisto/ |
| Bustools | Melsted et al.<sup>10</sup> | https://bustools.github.io |
| Cell Ranger | 10x Genomics | https://www.10xgenomics.com/support/software/cell-ranger/latest |
| Data processing and analysis code | This study | https://github.com/pachterlab/RMEJLBASBP_2024.git |
| Data processing and analysis Docker images | This study | https://hub.docker.com/repository/docker/josephrich98/scrnaseq_packages_and_versioning/general |

## RESOURCE AVAILABILITY

### Lead contact

Further information and requests for resources should be directed to and will be fulfilled by the lead contact, Lior Pachter.

### Materials availability

This study did not generate new unique reagents.

### Data and code availability

All original code has been deposited at GitHub and is publicly available as of the date of publication. The Docker image that provides the virtual environment in which all analysis was performed has been deposited at DockerHub and is publicly available as of the date of publication. Count matrix generation from FASTQ files was performed in a conda environment, also deposited on the GitHub repository. The original FASTQ files of the PBMC 10k dataset, shown in figures, and the PBMC 5k dataset, used for validation (figures not shown) can be found on the 10x Genomics website. The count matrices generated with kb-python and Cell Ranger from the PBMC 10k dataset FASTQ files have been deposited at Box and are publicly available as of the date of publication. Links to all materials can be found in the Key Resources table.

## METHOD DETAILS

### Count Matrix Generation

Count matrices were generated using either kb or Cell Ranger. The workflow used in most figures (unless stated otherwise) involved kb v0.28.0 (kallisto v0.50.1, bustools v0.43.1) and standard arguments. Other package versions used include Cell Ranger v7.2.0 and Cell Ranger v6.1.2. Reference transcriptomes for kb were built using kb ref with Ensembl release 111, downloaded with gget^40^; for Cell Ranger, the 2020-A reference transcriptome was used.

Read downsampling was simulated with the seqtk package. Cell downsampling was simulated with the “sample” function in base R.

### Analysis

All analysis was performed in R markdown files (R v4.3.1) with the reticulate package for Anaconda Python integration (v3.9.18)^41^. Data processing was performed with dplyr and tidyr^39^. All plots unless stated otherwise were generated with ggplot2^39^. The standard pipeline for Seurat v5.0.2^42^, and Scanpy v1.9.5^43^ were followed for comparison within and between packages.

### Filtering

Data were loaded and filtered by UMI counts with the Matrix and DropletUtils packages^44^. Filtering by cells and genes was performed with thresholding with standard values (minimum 3 cells per gene, minimum 200 genes per cell) for full-sized datasets. For datasets involving downsampled reads, these parameters were tuned to generally maximize the overlap of cell and gene sets between full and downsampled data. For datasets involving downsampled cells, the minimum cells per gene parameter was tuned to generally maximize the Jaccard index of gene sets, where Jaccard index is defined as intersection over union (i.e., the number of elements shared between two sets divided by the total number of elements between two combined sets). Filtering by mitochondrial gene content was performed with simple thresholding to remove cells with extremely high mitochondrial gene content as determined by violin plots (typically *>* 20% for the PBMC 10k dataset). Mitochondrial gene content was assessed by cross-referencing with the list of mitochondrial genes as determined by biomaRt^45^. Filtering results were visualized with UpSet plots using the UpSetR package^46^ (also used for all UpSet plots in this study) for the sets of cells and genes, and assessed using the Jaccard index between conditions.

### Normalization

Data normalization was performed by dividing each value in the count matrix by the total counts per cell, multiplying by a scaling factor of 10,000, and and then taking the log1p transform of this value. In other words, 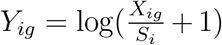 where 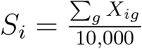. Normalization results were assessed by comparing the mean difference in entries after normalization of identical count matrices.

### Highly Variable Gene Selection

Highly variable genes were selected based on demonstrating high variance (normalized to mean expression) across all cells. In the case of Scanpy, subsequent analysis up until differential expression was performed exclusively on highly variable gene data. HVG set similarity was visualized with UpSet plots and assessed with Jaccard index.

### Principal Component Analysis

PCA analysis was performed, reducing the dimensionality of gene to the top 50 PCs. Analysis of PCA results was performed in three ways: by overlaying the PCA plots in two dimensions; by comparing the magnitudes of eigenvalues (Scree plots); and by comparing the sines of eigenvectors (where 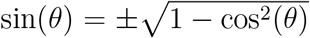 by the Pythagorean identity). The sine of the eigenvectors should be equal to 0 for identical vectors, and equal to 1 for perpendicular vectors, thus providing a quantification for similarity of vectors in high-dimensional space.

### Scaling and Regression

Gene expression data were scaled to have a mean of 0 and variance of 1 per gene across all cells, with user-defined clipping (upper-bound scaled expression). Optionally, features such as total counts and percentage of mitochondrial gene content could be regressed out during or after this step. Scaling and regression equivalence between packages was assessed by determining the equivalence of PCA embeddings when identical HVGs were fed into scaling and regression, and assessing the similarity of output through identical PCA steps.

### K-Nearest Neighbors and Shared Nearest Neighbors

The cell coordinates on the top 50 PCA embeddings were passed into KNN search algorithms in order to begin identifying similar groups of cells. For large datasets, Seurat used the Annoy algorithm through the RcppAnnoy package^26^, and Scanpy used NN Descent through the PyNNDescent package, called by umap-learn. The source code of Seurat’s FindNeighbors internal function calls was slightly modified in order to provide an interface to set the random seed of KNN graph generation with Annoy. SNN graphs were created from the KNN graphs based on neighborhood similarity of cells. The similarity of SNN graphs was assessed by computing the median Jaccard index and the median magnitude in the logarithm of the degree ratio between conditions.

The maximum Jaccard index *J*_*max*_ is equal to the degree ratio 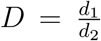, where *d*_1_ *< d*_2_. Jaccard index is intersection over union. The maximum overlap is given by *d*_1_, and the minimum union is given by *d*_2_. This ratio 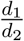 gives both the degree ratio and the upper bound on the Jaccard index.

### Clustering Analysis

Clustering was performed with the Louvain or Leiden algorithm. Seurat implemented Louvain itself, and Scanpy used louvain-igraph. Both packages used leidanalg for Leiden^47^. Assessment of cluster similarity was assessed visually with alluvial plots generated with the ggalluvial package^48^, and with the ARI calculated with the mclust package^49^.

The order of clusters and colors appearing in the alluvial plots was modified in a novel way from the capabilities of the ggalluvial package. The algorithm for the order of clusters, designed to minimize crossover in the alluvial plot and thus allow general cluster disorganization to correlate with agreement of clusters between groups, is as follows:

1. Order the left side from top down by decreasing cluster size
2. Order the right side such that it shares the nearest available rank to the cluster on the left with which is possesses its maximum Jaccard index
  a. If there are multiple clusters on the right that share the same cluster on the left for its highest Jaccard index, then these clusters are sorted by decreasing cluster size

Below is pseudocode that describes the implementation of this algorithm, which can be found implemented on this study’s GitHub repository^50^:

#### Algorithm 1 sort clusters by agreement

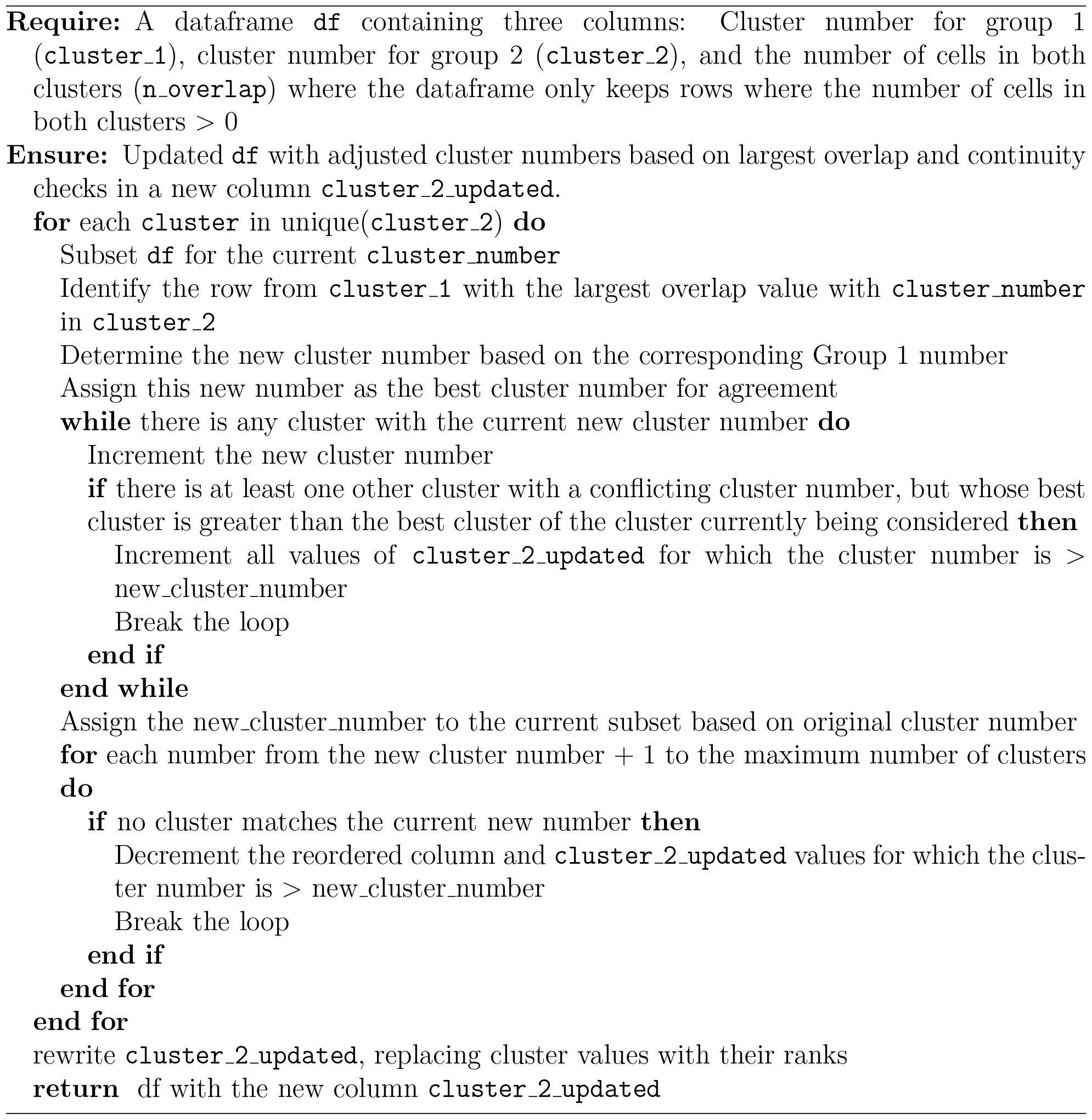

The algorithm for the order of colors follows the maximum weight matching algorithm in order to align the colors of the two groups as closely as possible.

### UMAP

UMAP plots were created from PCA-derived nearest neighbor graphs. By default, Seurat uses the uwot package^51^, and Scanpy uses umap-learn^29^. Analogous clusters were visually compared in local structure and relation to neighboring clusters to assess plot similarity.

UMAP data extended analysis was performed by constructing a KNN graph from the UMAP space, passing this KNN graph through Leiden clustering, and further projecting the UMAP-derived KNN graph in UMAP space. KNN construction from UMAP space was performed with an exact KNN graph implementation from the FNN package, which implements an exact KNN with Kd trees^52^. Leiden clustering was implemented with the bluster package^53^, which internally implements an exact KNN algorithm for input into Leiden using the kmeans k-nearest neighbor algorithm^54^. Further UMAP projection of the FNN-derived KNN graphs was implemented with the uwot package (the same underlying package as Seurat).

### Differential Expression

DE analysis was performed on the clustered genes by computing a logFC and p-value for each gene per cluster compared to all other clusters. Optionally, genes per cluster could be filtered with thresholding for logFC, p-value, or percentage of cells expressing the gene in a reference group to only select the best markers. We wrote a function to convert the data structure holding the values of percentage of cells in each reference group expressing a gene from a pandas dataframe to a numpy recarray to make it compatible with the other DE statistics stored in the Scanpy Anndata object. p-values were adjusted with the Bonferroni or Benjamini-Hochberg correction. Analysis of significant marker gene similarity (all genes across all clusters with adjusted p-value *<* 0.05) and marker similarity (all cluster-specific marker genes after any implemented filtering) was performed with UpSet plots and Jaccard indices of gene sets. Analysis of logFC similarity was performed with scatterplots, computation of the CCC, median and mean magnitude of difference of logFC for analogous marker genes, and a PCA best-fit line. CCC is defined as follows:

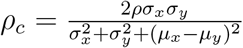

Analysis of adjusted p-value similarity was assessed with log scatterplots and by computing the percentage of marker genes which flipped across the significance threshold of p=0.05.

## QUANTIFICATION AND STATISTICAL ANALYSIS

Data were loaded and filtered by UMI counts with the Matrix package^44^. Each pipeline generally consisted of the following steps: filtering (UMIs, cells and genes, mitochondrial gene content), log normalization, HVG selection, feature regression, scaling, PCA, KNN and SNN graph formation, clustering, UMAP projection, and DE analysis (Supplemental Figure 1). Mitochondrial gene content was facilitated with biomaRt^45^. Filtering similarity was assessed by comparing similarity of cell and gene sets with UpSet plots. Normalization similarity between packages was assessed by providing the same data to each normalization function as input and comparing the similarity of output matrix values. HVG selection similarity was assessed by comparing overlap with UpSet plots and Jaccard index. PCA similarity was assessed by overlaying the PCA plots on the first two PCs, comparing the difference in corresponding eigenvector values on Scree plots, and comparing the sine of corresponding eigenvectors. SNN graph similarity was assessed by computing the Jaccard index for the overlap in neighborhoods, as well as the median magnitude in the ratio in degree of neighborhoods, for each cell between conditions. Cluster similarity was assessed by visually determining the degree of alignment in alluvial plots between clusters, as well as computing the ARI. UMAP similarity was determined by visually comparing plots, as well as by computing KNN graphs from the UMAP space, followed by subsequent Leiden clustering and UMAP projection of these UMAP-space derived KNN graphs, with assessment completed as before (Jaccard indices of KNN graph neighborhoods, alluvial plotting and ARI computation for clusters, and visual comparison of UMAP plots). DE similarity was assessed with comparing overlap of the union of significant marker genes across all clusters between conditions with UpSet plots. When cluster assignments could be unified between conditions, additional analysis of DE similarity was performed for each marker per cluster by calculating similarity of marker genes per cluster, by plotting logFC and computing the CCC and PCA fit line, and by plotting adjusted p-value between conditions and computing the percentage of markers which flipped across the p=0.05 threshold.

