## Supplemental Information for "The impact of package selection and versioning on single-cell RNA-seq analysis"

April 9, 2024

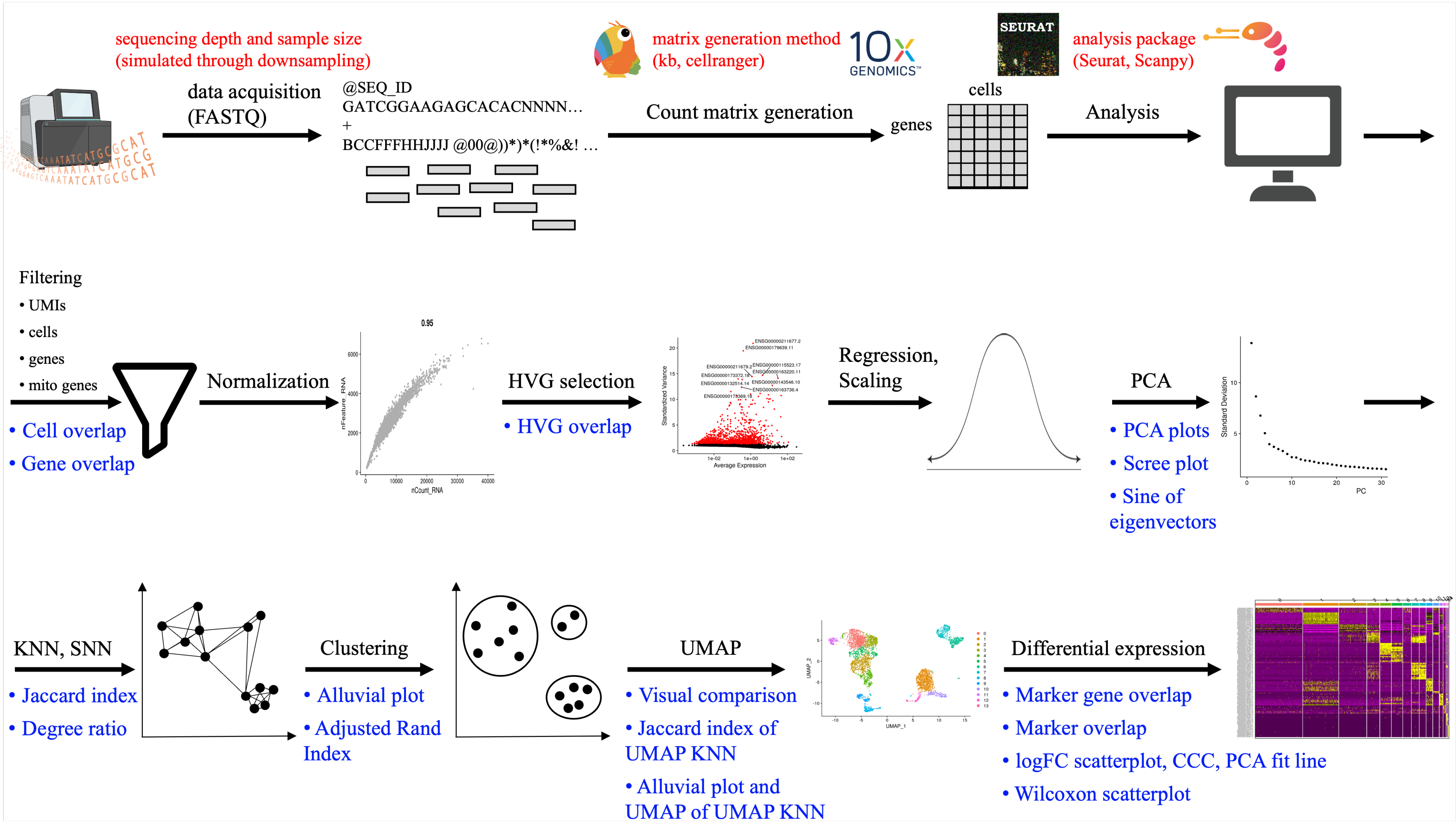

sequencing depth and sample size
(simulated through downsampling)
data acquisition
(FASTQ)
@SEQ_ID
GATCGGAAGAGCACACNNNN…
+
BCCFFFHHJJJJ @00@))*)*(!*%&! …
matrix generation method
(kb, cellranger)
10x GENOMICS
Count matrix generation
SEURAT
analysis package
(Seurat, Scanpy)
cells
genes
Analysis
Filtering
• UMIs
• cells
• genes
• mito genes
• Cell overlap
• Gene overlap
Normalization
HVG selection
• HVG overlap
Regression, Scaling
PCA
• PCA plots
• Scree plot
• Sine of eigenvectors
KNN, SNN
• Jaccard index
• Degree ratio
Clustering
• Alluvial plot
• Adjusted Rand Index
UMAP
• Visual comparison
• Jaccard index of UMAP KNN
• Alluvial plot and UMAP of UMAP KNN
Differential expression
• Marker gene overlap
• Marker overlap
• logFC scatterplot, CCC, PCA fit line
• Wilcoxon scatterplot

**Supplemental Figure 1:** scRNA-seq matrix generation and analysis overview. Red = sources of variability explored in this study; Blue = plots and metrics used to assess function similarity

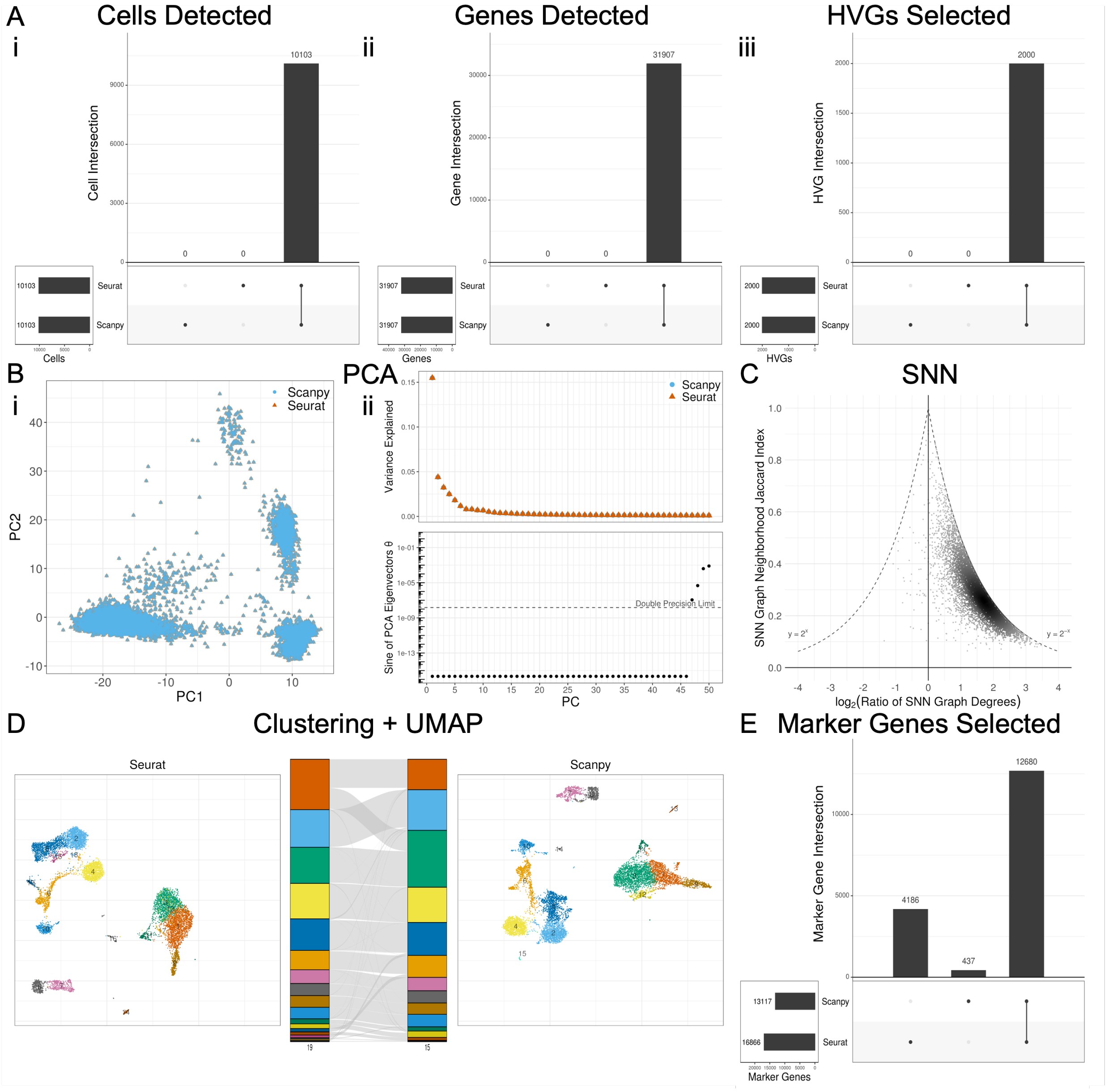


**Supplemental Figure 2:** Seurat vs. Scanpy, aligned function arguments (Seurat-like), sequential analysis. (A) Filtering and HVG selection analysis UpSet plots consisting of overlap of sets of cells, genes, and HVGs. (B) PCA analysis through projection onto first 2 PCs, Scree plot eigenvalue comparison, and sine of eigenvectors. Black lines = point mapping between conditions. (C) KNN/SNN analysis through SNN neighborhood Jaccard index and degree ratio (Seurat/Scanpy) per cell. (D) Clustering and UMAP analysis through UMAP plots of each condition, with alluvial plot showing cluster assignment mapping and degree of agreement. (E) Differential expression analysis UpSet plot through overlap of all significant ($p < 0.05$) marker genes across all clusters.

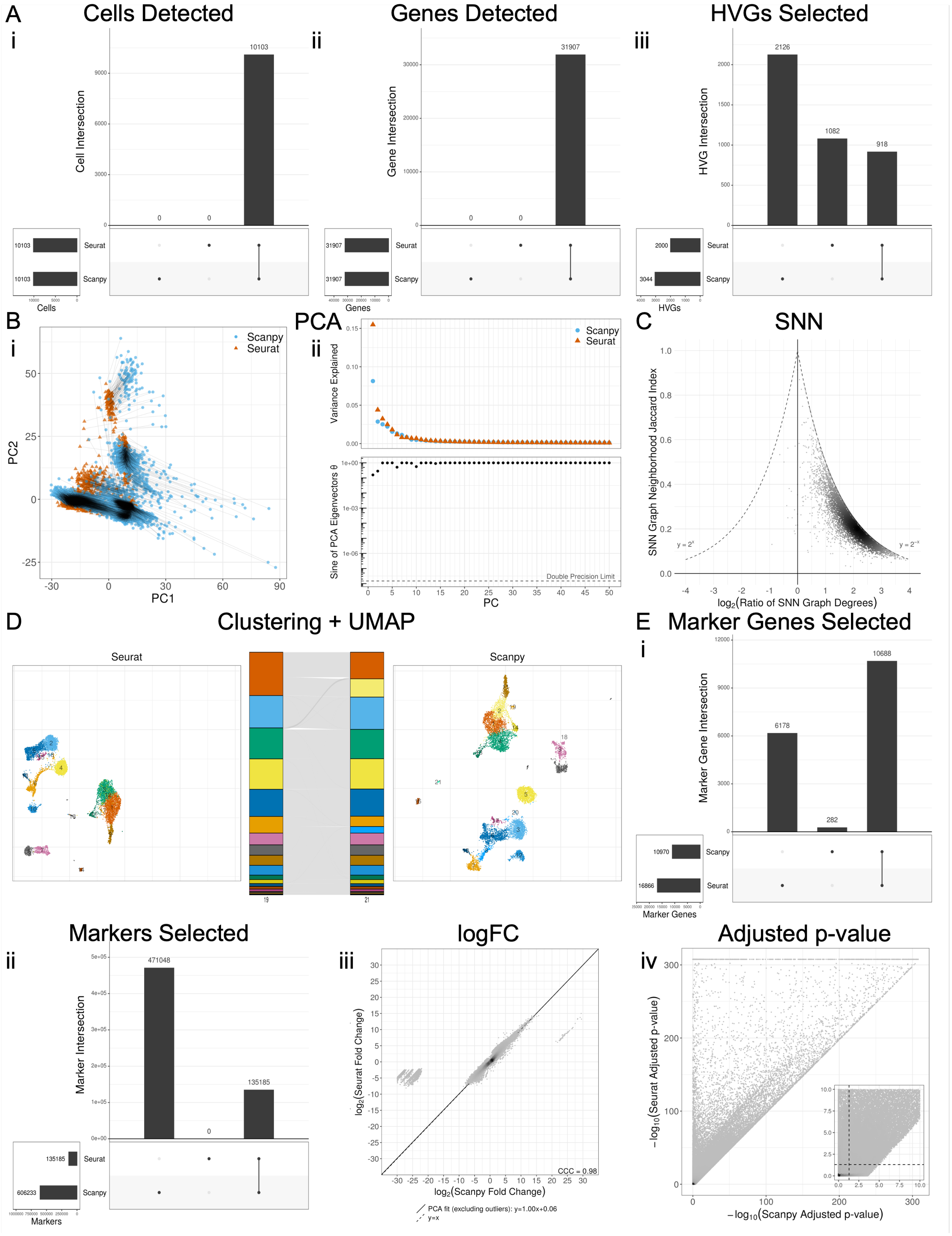
A
Cells Detected
Genes Detected
HVGs Selected
Cell Intersection
Gene Intersection
HVG Intersection
Seurat
Scanpy
Cells
Genes
HVGs
B
PCA
PC1
PC2
Variance Explained
Sine of PCA Eigenvectors θ
Double Precision Limit
PC
C
SNN
SNN Graph Neighborhood Jaccard Index
log2(Ratio of SNN Graph Degrees)
D
Clustering + UMAP
Seurat
Scanpy
E
Marker Genes Selected
Marker Gene Intersection
Marker Genes
Markers Selected
Marker Intersection
Markers
logFC
log2(Seurat Fold Change)
log2(Scanpy Fold Change)
CCC = 0.98
PCA fit (excluding outliers): y=1.00x+0.06
y=x
Adjusted p-value
−log10(Seurat Adjusted p-value)
−log10(Scanpy Adjusted p-value)

**Supplemental Figure 3:** Seurat vs. Scanpy, default function arguments, controlled analysis. (A) Filtering and HVG selection analysis UpSet plots consisting of overlap of sets of cells, genes, and HVGs. (B) PCA analysis through projection onto first 2 PCs, Scree plot eigenvalue comparison, and sine of eigenvectors. Black lines = point mapping between conditions. (C) KNN/SNN analysis through SNN neighborhood Jaccard index and degree ratio (Seurat/Scanpy) per cell. (D) Clustering and UMAP analysis through UMAP plots of each condition, with alluvial plot showing cluster assignment mapping and degree of agreement. (E) Differential expression analysis through overlap of all significant ($p < 0.05$) marker genes across all clusters. (F) Differential expression analysis with aligned clusters of overlap of all markers. (G) Differential expression analysis with aligned clusters of logFC similarity. (H) Differential expression analysis with aligned clusters of adjusted p-value similarity.

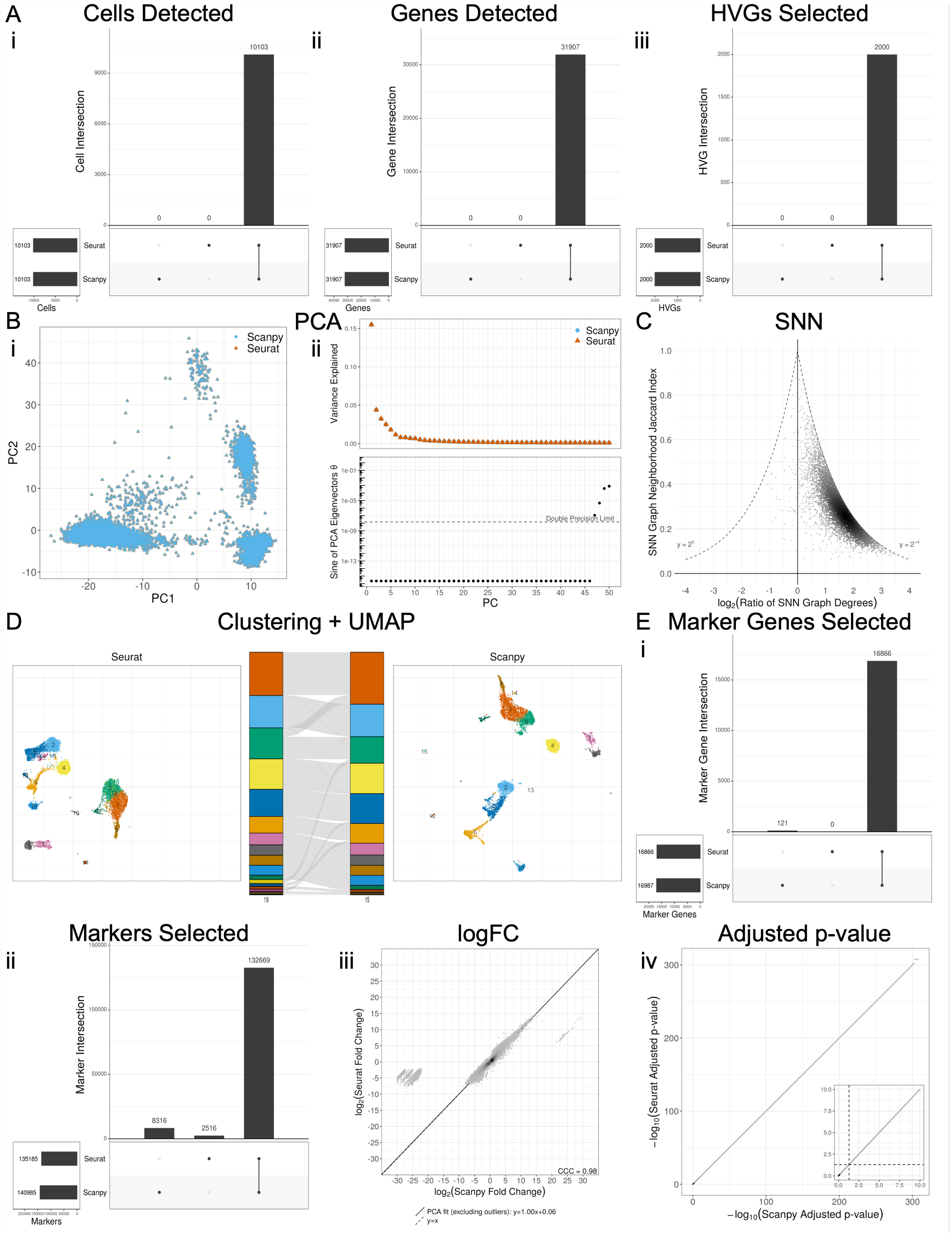

A
Cells Detected
i
Cell Intersection
10103
0
0
Seurat
Scanpy
Cells
Genes Detected
ii
Gene Intersection
31907
Genes
HVGs Selected
iii
HVG Intersection
2000
HVGs
B
i
Scanpy
Seurat
PC2
PC1
PCA
ii
Variance Explained
Sine of PCA Eigenvectors θ
Double Precision Limit
PC
C
SNN
SNN Graph Neighborhood Jaccard Index
log2(Ratio of SNN Graph Degrees)
y = 2^x
y = 2^-x
D
Clustering + UMAP
Seurat
Scanpy
E
Marker Genes Selected
i
Marker Gene Intersection
16866
121
Marker Genes
ii
Markers Selected
Marker Intersection
132669
8316
2516
Markers
iii
logFC
log2(Seurat Fold Change)
log2(Scanpy Fold Change)
CCC = 0.98
PCA fit (excluding outliers): y=1.00x+0.06
y=x
iv
Adjusted p-value
−log10(Seurat Adjusted p-value)
−log10(Scanpy Adjusted p-value)

**Supplemental Figure 4:** Seurat vs. Scanpy, aligned function arguments (Seurat-like), controlled analysis. (A) Filtering and HVG selection analysis UpSet plots consisting of overlap of sets of cells, genes, and HVGs. (B) PCA analysis through projection onto first 2 PCs, Scree plot eigenvalue comparison, and sine of eigenvectors. Black lines = point mapping between conditions. (C) KNN/SNN analysis through SNN neighborhood Jaccard index and degree ratio (Seurat/Scanpy) per cell. (D) Clustering and UMAP analysis through UMAP plots of each condition, with alluvial plot showing cluster assignment mapping and degree of agreement. (E) Differential expression analysis through overlap of all significant ($p < 0.05$) marker genes across all clusters. (F) Differential expression analysis with aligned clusters of overlap of all markers. (G) Differential expression analysis with aligned clusters of logFC similarity. (H) Differential expression analysis with aligned clusters of adjusted p-value similarity.

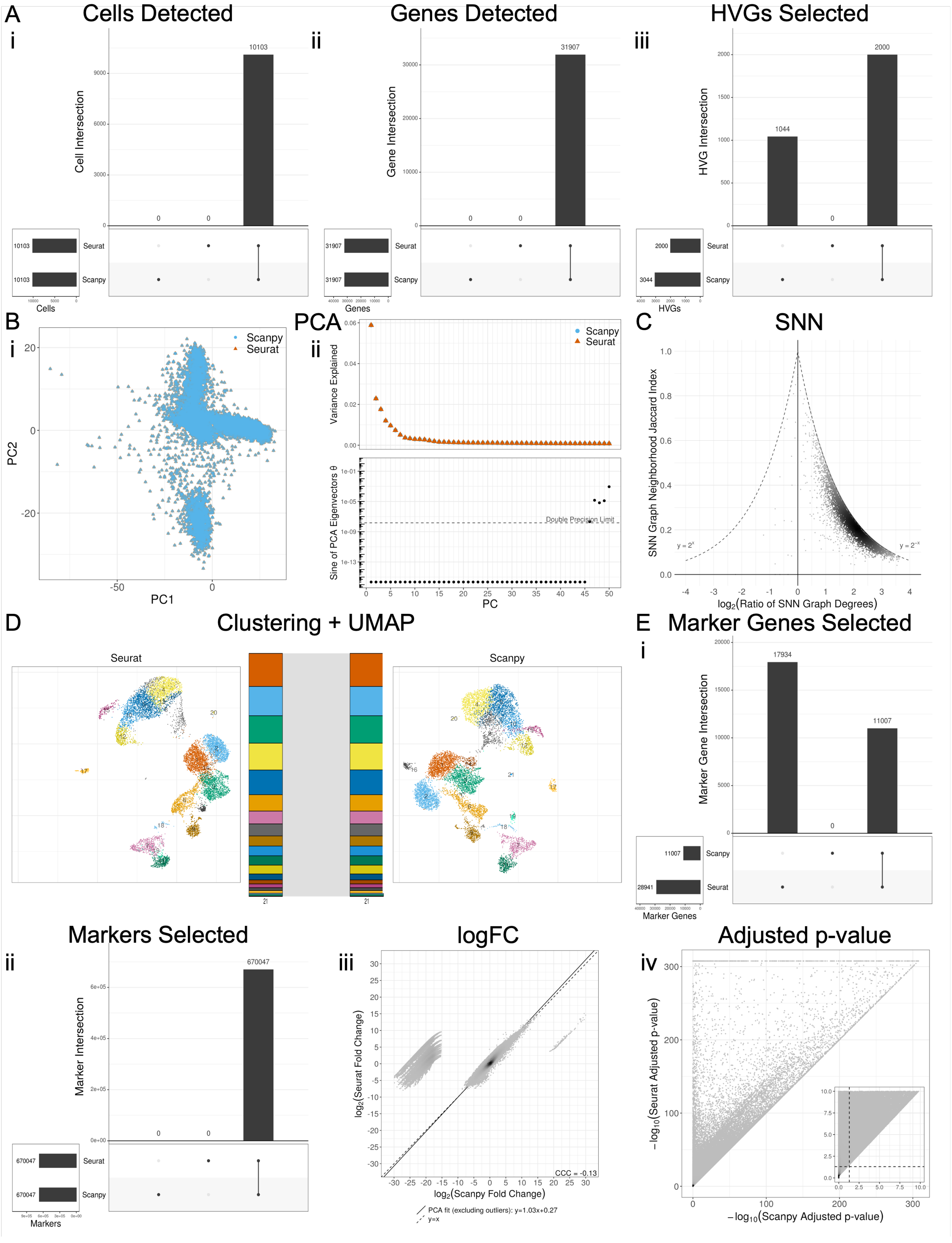

A
Cells Detected
i
Cell Intersection
Cells
Seurat
Scanpy
Genes Detected
ii
Gene Intersection
Genes
HVGs Selected
iii
HVG Intersection
HVGs
B
i
Scanpy
Seurat
PC1
PC2
PCA
ii
Variance Explained
Sine of PCA Eigenvectors θ
Double Precision Limit
PC
C
SNN
SNN Graph Neighborhood Jaccard Index
log2(Ratio of SNN Graph Degrees)
D
Clustering + UMAP
E
Marker Genes Selected
Marker Gene Intersection
Marker Genes
Markers Selected
Marker Intersection
Markers
logFC
log2(Seurat Fold Change)
log2(Scanpy Fold Change)
CCC = -0.13
PCA fit (excluding outliers): y=1.03x+0.27
y=x
Adjusted p-value
iv
−log10(Seurat Adjusted p-value)
−log10(Scanpy Adjusted p-value)

**Supplemental Figure 5:** Seurat vs. Scanpy, aligned function arguments (Scanpy-like), controlled analysis. (A) Filtering and HVG selection analysis UpSet plots consisting of overlap of sets of cells, genes, and HVGs. (B) PCA analysis through projection onto first 2 PCs, Scree plot eigenvalue comparison, and sine of eigenvectors. Black lines = point mapping between conditions. (C) KNN/SNN analysis through SNN neighborhood Jaccard index and degree ratio (Seurat/Scanpy) per cell. (D) Clustering and UMAP analysis through UMAP plots of each condition, with alluvial plot showing cluster assignment mapping and degree of agreement. (E) Differential expression analysis through overlap of all significant ($p < 0.05$) marker genes across all clusters. (F) Differential expression analysis with aligned clusters of overlap of all markers. (G) Differential expression analysis with aligned clusters of logFC similarity. (H) Differential expression analysis with aligned clusters of adjusted p-value similarity.

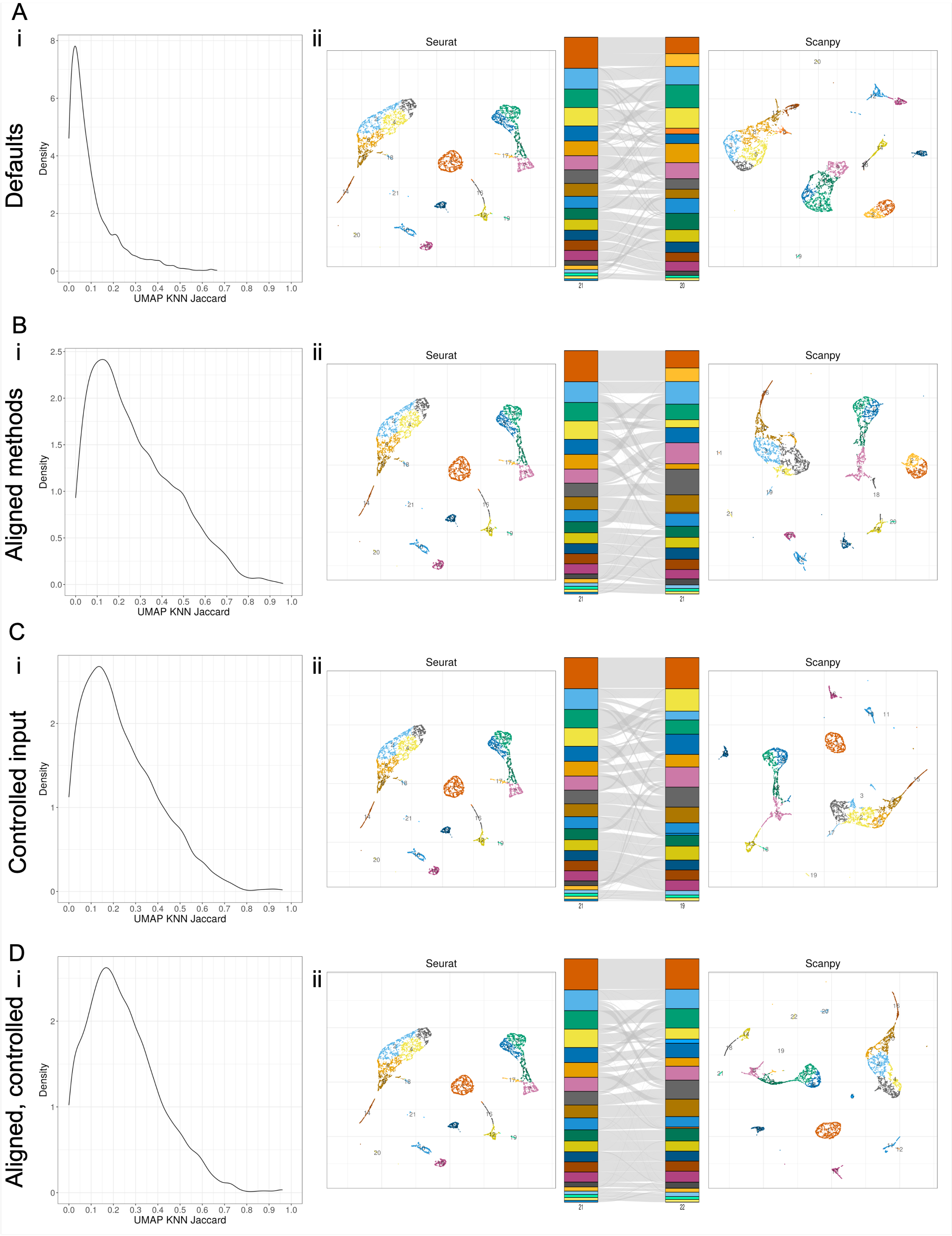
A
i
Defaults
Density
UMAP KNN Jaccard
ii
Seurat
Scanpy
B
i
Aligned methods
Density
UMAP KNN Jaccard
ii
Seurat
Scanpy
C
i
Controlled input
Density
UMAP KNN Jaccard
ii
Seurat
Scanpy
D
i
Aligned, controlled
Density
UMAP KNN Jaccard
ii
Seurat
Scanpy

**Supplemental Figure 6:** Extended UMAP analysis for Seurat vs. Scanpy showing density plot of UMAP-derived KNN graph neighborhood Jaccard indices, alluvial plot of leiden clustering performed on UMAP-derived KNN graph, and UMAP projection of UMAP-derived KNN graph colored with leiden clustering results. (A) Default methods, sequential analysis. (B) Aligned methods, sequential analysis. (C) Default methods, controlled analysis. (D) Aligned methods, controlled analysis.

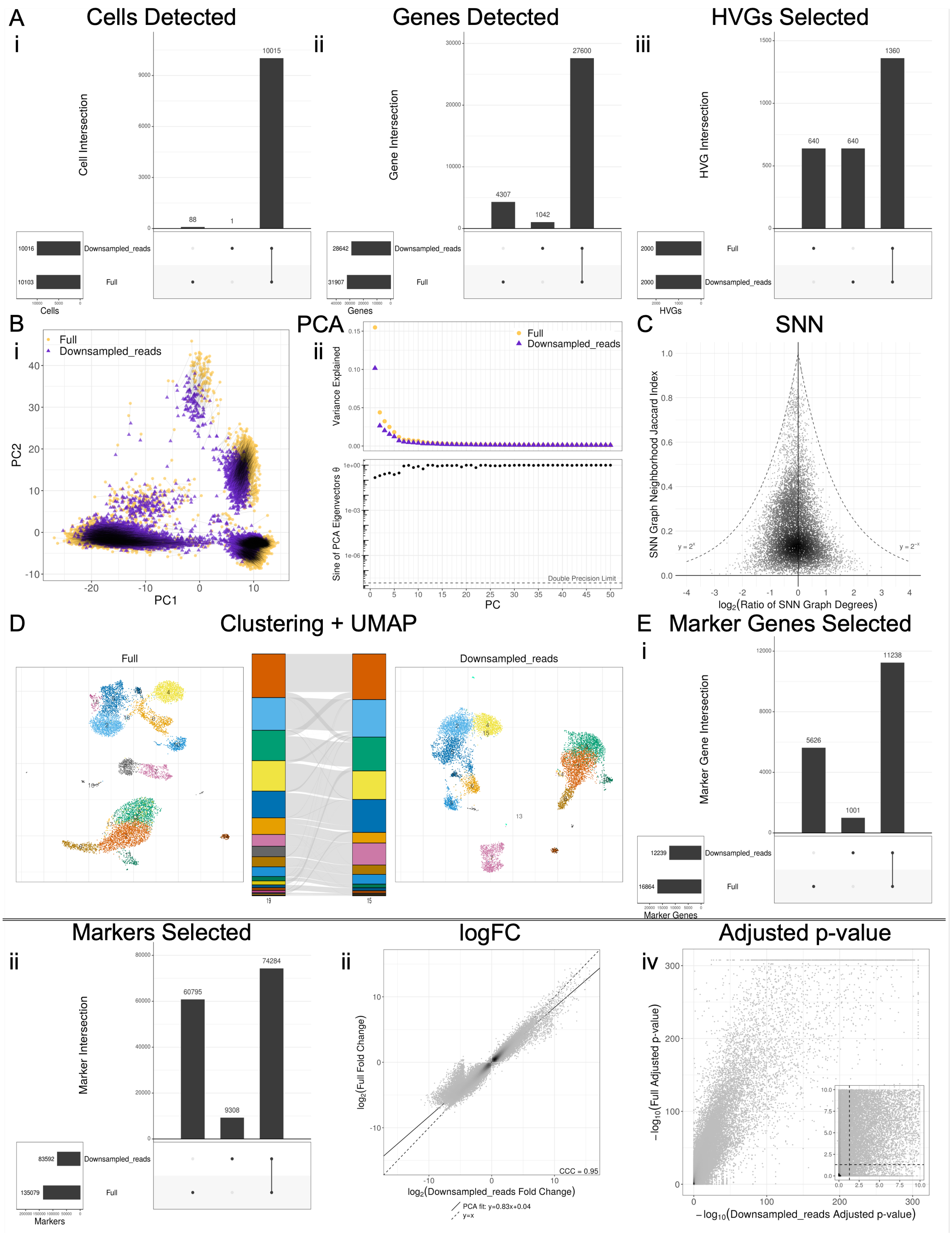

A
Cells Detected
i
Cell Intersection
10015
88
1
Downsampled_reads
Full
10016
10103
Cells
ii
Genes Detected
Gene Intersection
27600
4307
1042
28642
31907
Genes
iii
HVGs Selected
HVG Intersection
1360
640
640
2000
2000
HVGs
B
i
Full
Downsampled_reads
PC1
PC2
PCA
ii
Variance Explained
Sine of PCA Eigenvectors θ
Double Precision Limit
PC
C
SNN
SNN Graph Neighborhood Jaccard Index
log2(Ratio of SNN Graph Degrees)
D
Clustering + UMAP
E
Marker Genes Selected
Marker Gene Intersection
11238
5626
1001
12239
16864
Marker Genes
Markers Selected
Marker Intersection
74284
60795
9308
83592
135079
Markers
logFC
log2(Full Fold Change)
log2(Downsampled_reads Fold Change)
CCC = 0.95
PCA fit: y=0.83x+0.04
y=x
Adjusted p-value
iv
−log10(Full Adjusted p-value)
−log10(Downsampled_reads Adjusted p-value)

**Supplemental Figure 7:** Read downsampling to 4% of original size vs. full dataset in Seurat, with a fraction that displays variation comparable to Seurat vs. Scanpy. (A) Filtering and HVG selection analysis UpSet plots consisting of overlap of sets of cells, genes, and HVGs. (B) PCA analysis through projection onto first 2 PCs, Scree plot eigenvalue comparison, and sine of eigenvectors. Black lines = point mapping between conditions. (C) KNN/SNN analysis through SNN neighborhood Jaccard index and degree ratio (full/downsampled) per cell. (D) Clustering and UMAP analysis through UMAP plots of each condition, with alluvial plot showing cluster assignment mapping and degree of agreement. (E) Differential expression analysis through overlap of all significant ($p < 0.05$) marker genes across all clusters. (F) Differential expression analysis with aligned clusters of overlap of all markers. (G) Differential expression analysis with aligned clusters of logFC similarity. (H) Differential expression analysis with aligned clusters of adjusted p-value similarity.

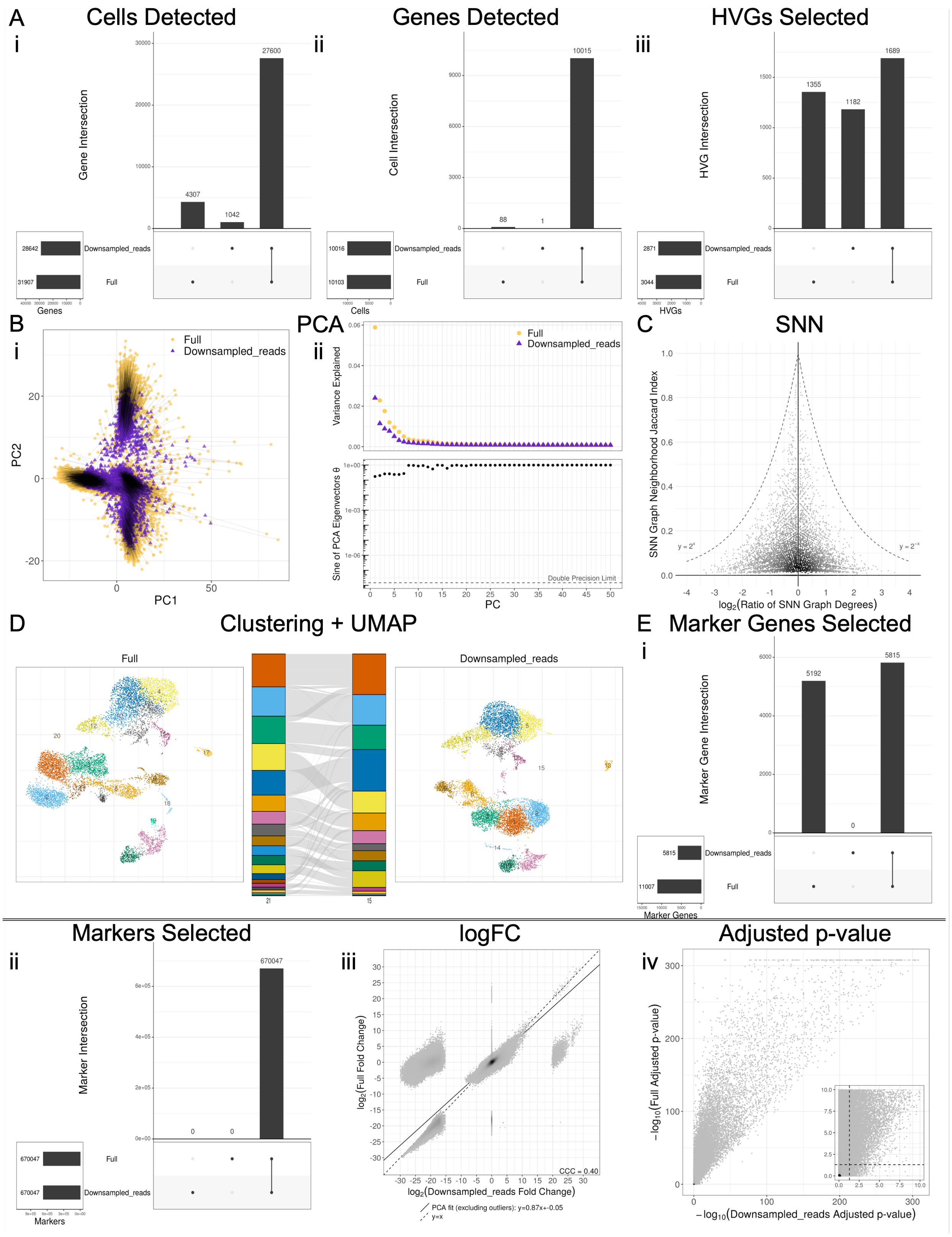

A
Cells Detected
Genes Detected
HVGs Selected
i
ii
iii
Gene Intersection
Cell Intersection
HVG Intersection
27600
4307
1042
10015
88
1
1689
1355
1182
Downsampled_reads
Full
28642
31907
10016
10103
2871
3044
Genes
Cells
HVGs
B
PCA
C
SNN
Full
Downsampled_reads
PC2
PC1
Variance Explained
Sine of PCA Eigenvectors θ
Double Precision Limit
PC
SNN Graph Neighborhood Jaccard Index
$\log_2$(Ratio of SNN Graph Degrees)
$y = 2^x$
$y = 2^{-x}$
D
Clustering + UMAP
Full
Downsampled_reads
E
Marker Genes Selected
Marker Gene Intersection
5192
0
5815
11007
Marker Genes
Markers Selected
logFC
Adjusted p-value
iv
Marker Intersection
670047
0
Markers
$\log_2$(Full Fold Change)
$\log_2$(Downsampled_reads Fold Change)
CCC = 0.40
PCA fit (excluding outliers): y=0.87x+-0.05
y=x
$-\log_{10}$(Full Adjusted p-value)
$-\log_{10}$(Downsampled_reads Adjusted p-value)

**Supplemental Figure 8:** Read downsampling to 4% of original size vs. full dataset in Scanpy, with a fraction that displays variation comparable to Seurat vs. Scanpy. (A) Filtering and HVG selection analysis UpSet plots consisting of overlap of sets of cells, genes, and HVGs. (B) PCA analysis through projection onto first 2 PCs, Scree plot eigenvalue comparison, and sine of eigenvectors. Black lines = point mapping between conditions. (C) KNN/SNN analysis through SNN neighborhood Jaccard index and degree ratio (full/-downsampled) per cell. (D) Clustering and UMAP analysis through UMAP plots of each condition, with alluvial plot showing cluster assignment mapping and degree of agreement. (E) Differential expression analysis through overlap of all significant ($p < 0.05$) marker genes across all clusters. (F) Differential expression analysis with aligned clusters of overlap of all markers. (G) Differential expression analysis with aligned clusters of logFC similarity. (H) Differential expression analysis with aligned clusters of adjusted p-value similarity.

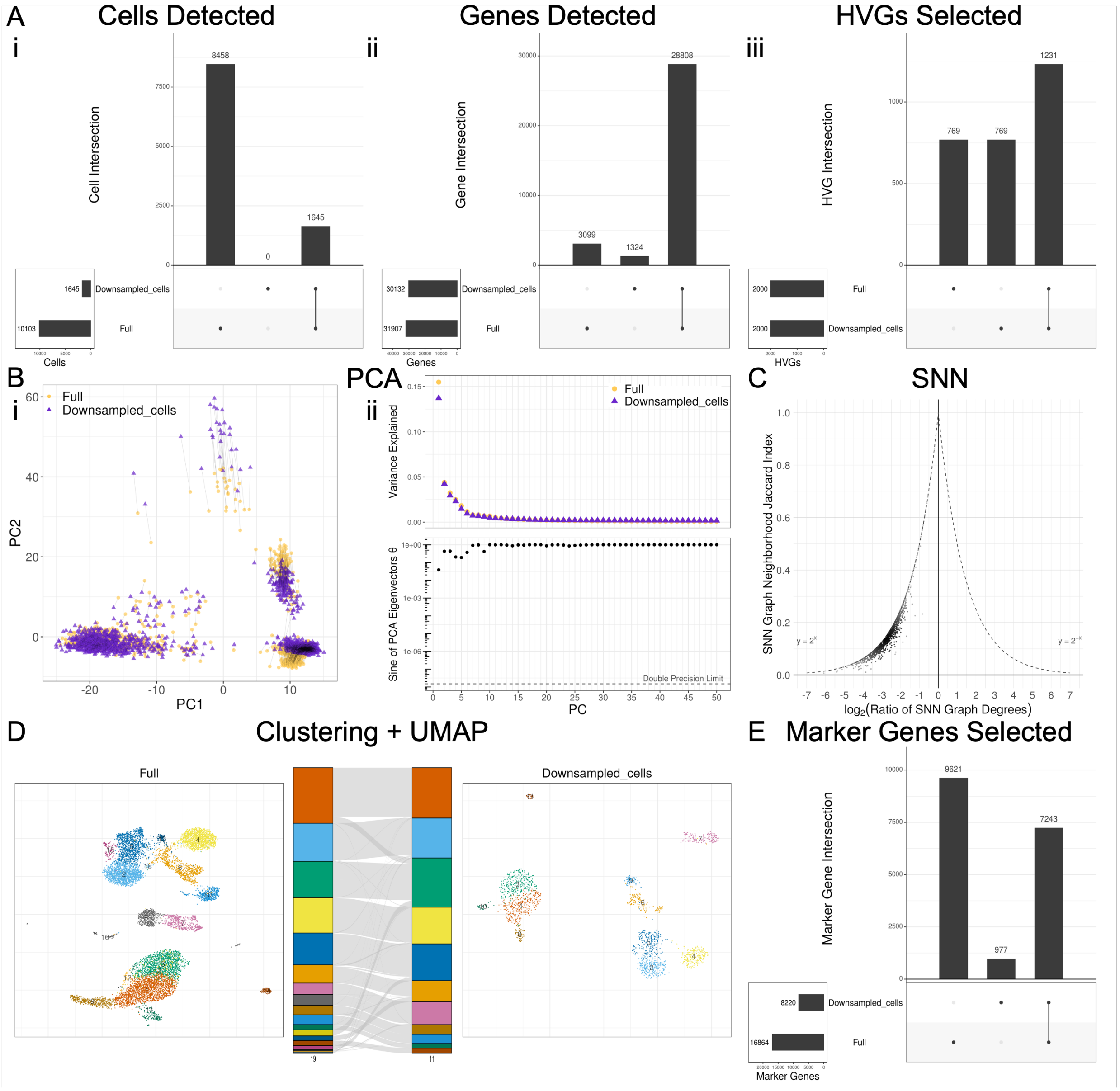


**Supplemental Figure 9:** Cell downsampling to 16% of original size vs. full dataset in Seurat, with a fraction that displays variation comparable to Seurat vs. Scanpy. (A) Filtering and HVG selection analysis UpSet plots consisting of overlap of sets of cells, genes, and HVGs. (B) PCA analysis through projection onto first 2 PCs, Scree plot eigenvalue comparison, and sine of eigenvectors. Black lines = point mapping between conditions. (C) KNN/SNN analysis through SNN neighborhood Jaccard index and degree ratio (full/downsampled) per cell. (D) Clustering and UMAP analysis through UMAP plots of each condition, with alluvial plot showing cluster assignment mapping and degree of agreement. (E) Differential expression analysis UpSet plot through overlap of all significant ($p < 0.05$) marker genes across all clusters.

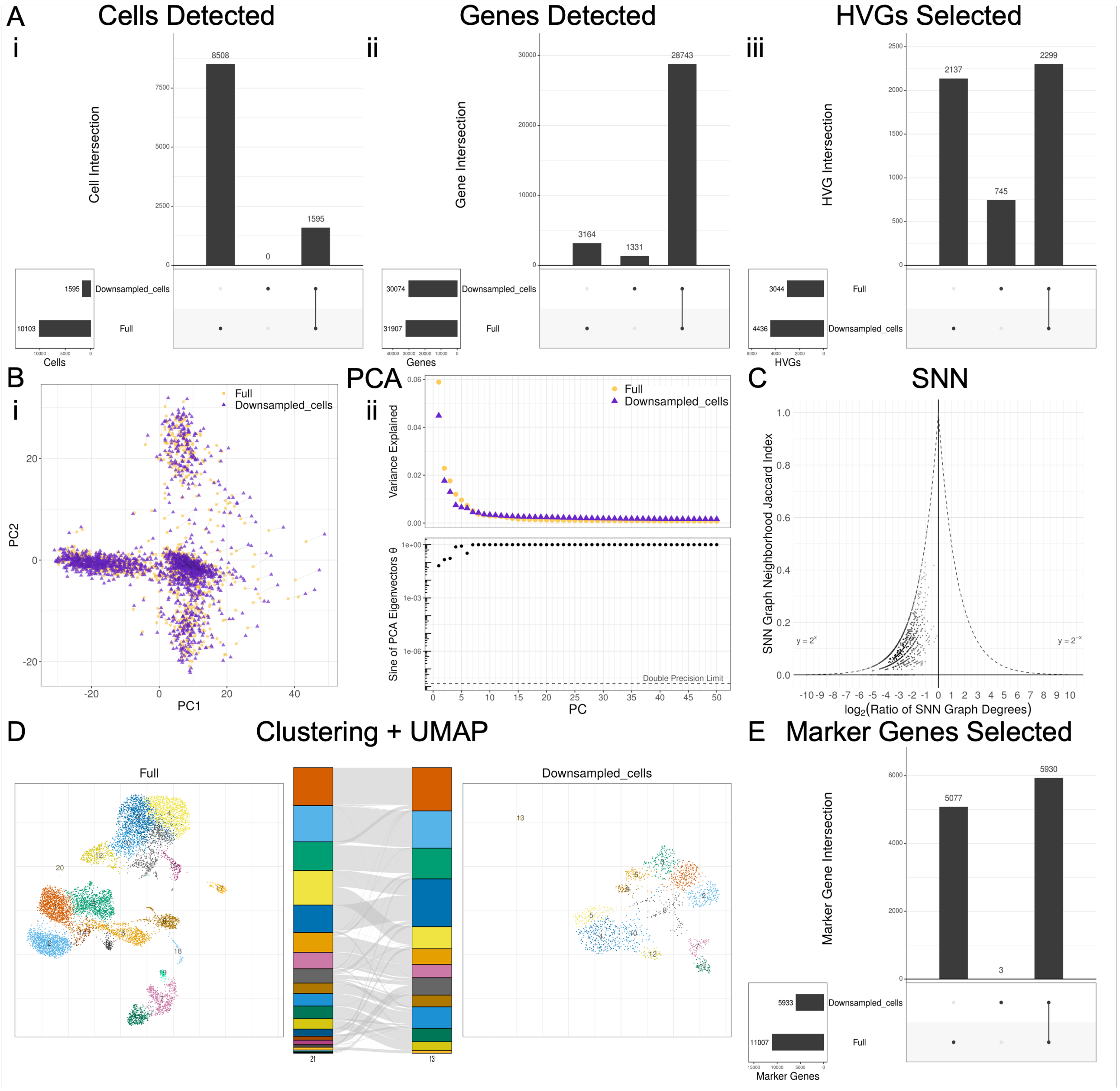


**Supplemental Figure 10:** Cell downsampling to 16% of original size vs. full dataset in Scanpy, with a fraction that displays variation comparable to Seurat vs. Scanpy. (A) Filtering and HVG selection analysis UpSet plots consisting of overlap of sets of cells, genes, and HVGs. (B) PCA analysis through projection onto first 2 PCs, Scree plot eigenvalue comparison, and sine of eigenvectors. Black lines = point mapping between conditions. (C) KNN/SNN analysis through SNN neighborhood Jaccard index and degree ratio (full/-downsampled) per cell. (D) Clustering and UMAP analysis through UMAP plots of each condition, with alluvial plot showing cluster assignment mapping and degree of agreement. (E) Differential expression analysis UpSet plot through overlap of all significant ($p < 0.05$) marker genes across all clusters.

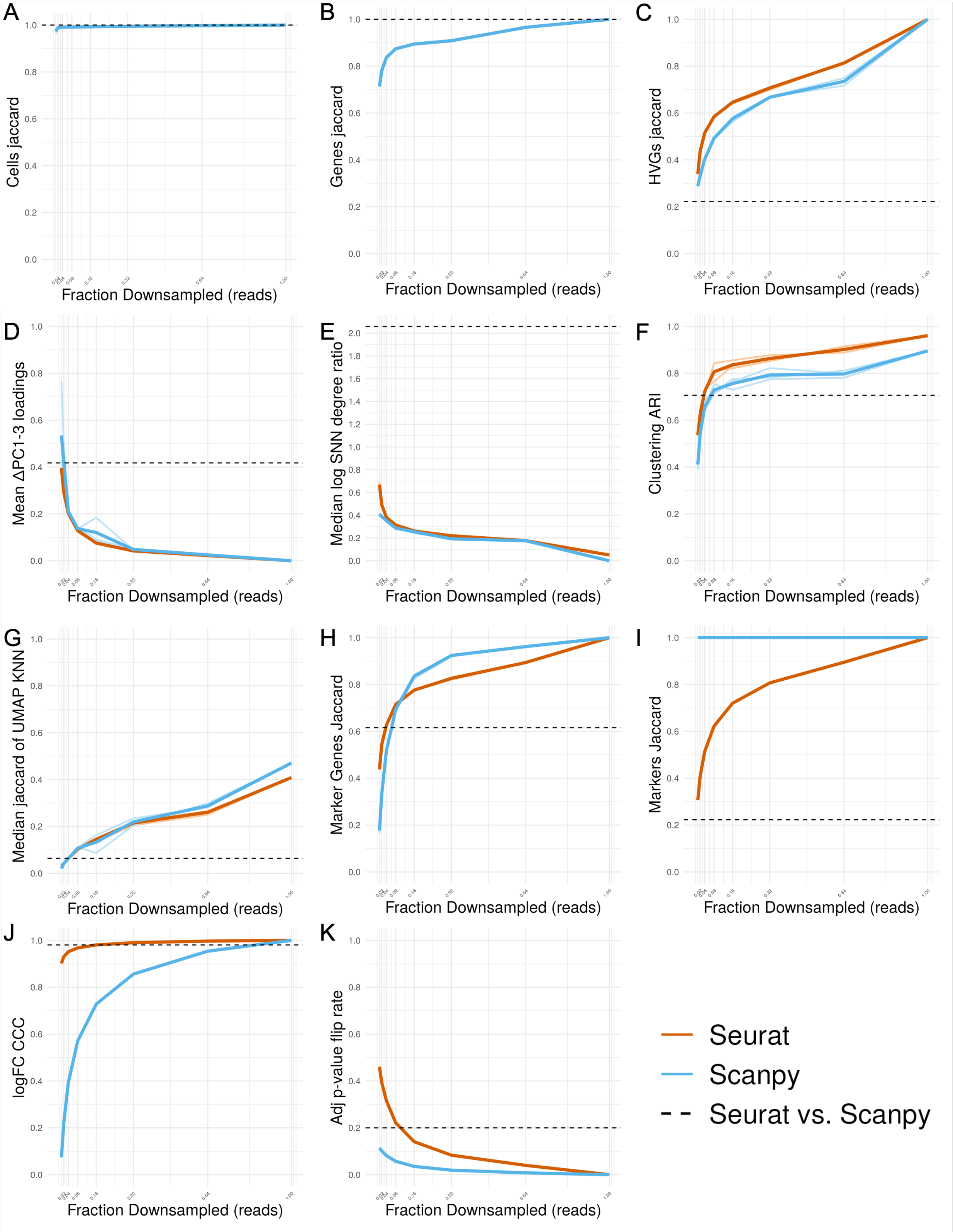

A
Cells jaccard
B
Genes jaccard
C
HVGs jaccard
D
Mean ΔPC1-3 loadings
E
Median log SNN degree ratio
F
Clustering ARI
G
Median jaccard of UMAP KNN
H
Marker Genes Jaccard
I
Markers Jaccard
J
logFC CCC
K
Adj p-value flip rate
Fraction Downsampled (reads)
Seurat
Scanpy
Seurat vs. Scanpy

**Supplemental Figure 11:** Metrics vs. fraction of reads downsampled. (A) Jaccard index of cell overlap. (B) Jaccard index of gene overlap. (C) Jaccard index of HVG overlap. (D) Mean of difference in corresponding PC loadings for PCs 1-3. (E) Median log(SNN degree ratio). (F) Adjusted Rand Index of clusters. (G) Median Jaccard index of UMAP-derived KNN graphs for each cell. (H) Jaccard index of sets of significant ($p < 0.05$) marker genes across all clusters. (I) Jaccard index of all markers. (J) CCC of logFC scatterploz. (K) Fraction of adjusted p-value that flipped across p=0.05 threshold. Orange = Seurat; Blue = Scanpy; lighter lines = replicates of downsampling across random seeds; dashed line = value recorded by Seurat vs. Scanpy with default settings.

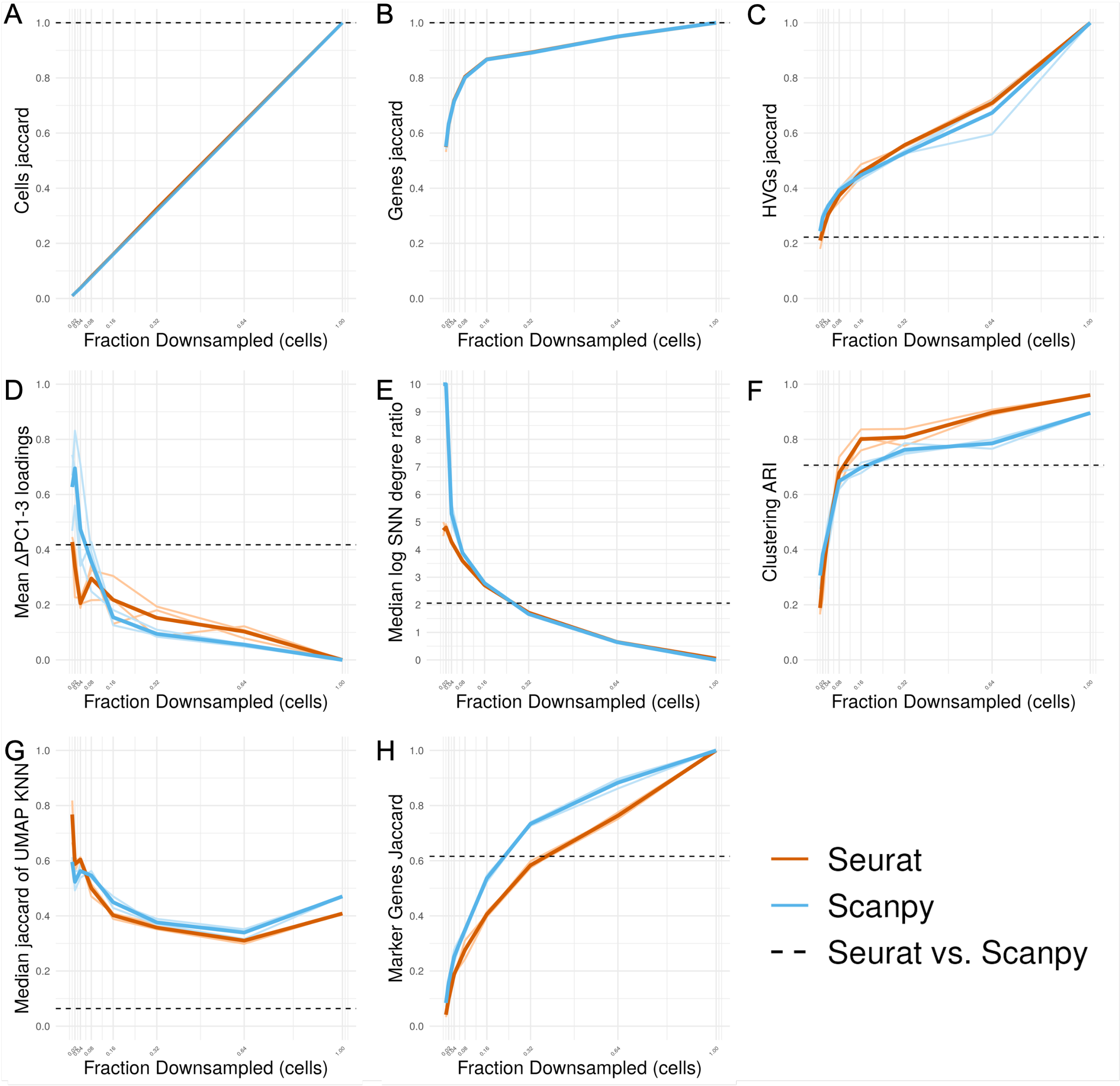


**Supplemental Figure 12:** Metrics vs. fraction of cells downsampled. (A) Jaccard index of cell overlap. (B) Jaccard index of gene overlap. (C) Jaccard index of HVG overlap. (D) Mean of difference in corresponding PC loadings for PCs 1-3. (E) Median log(SNN degree ratio). (F) Adjusted Rand Index of clusters. (G) Median Jaccard index of UMAP-derived KNN graphs for each cell. (H) Jaccard index of sets of significant ($p < 0.05$) marker genes across all clusters. (I) Jaccard index of all markers. (J) CCC of logFC scatterplot. (K) Fraction of adjusted p-value that flipped across p=0.05 threshold. Orange = Seurat; Blue = Scanpy; lighter lines = replicates of downsampling across random seeds; dashed line = value recorded by Seurat vs. Scanpy with default settings.

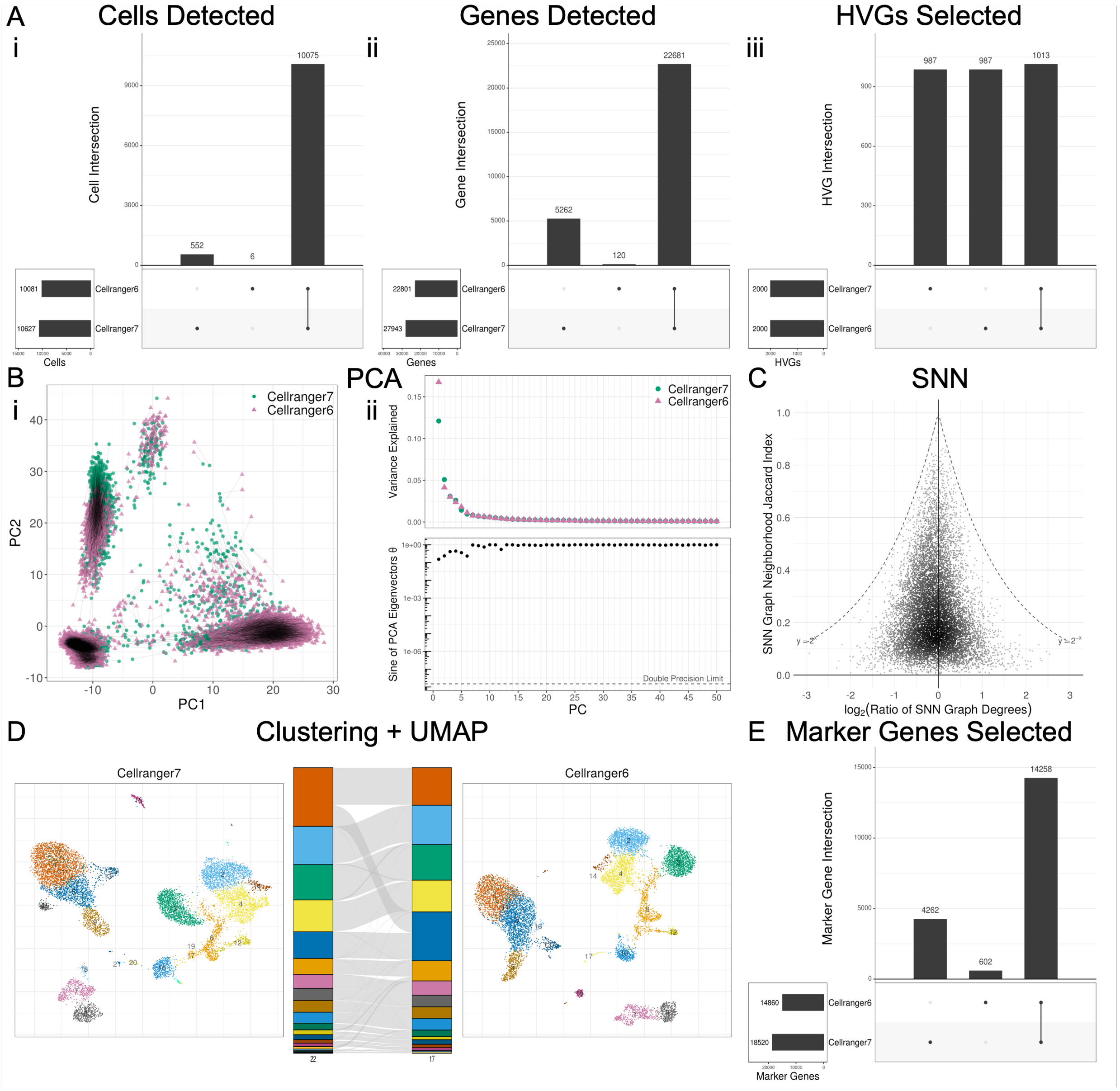


**Supplemental Figure 13:** Cell Ranger version control (v7 vs. v6). (A) Filtering and HVG selection analysis UpSet plots consisting of overlap of sets of cells, genes, and HVGs. (B) PCA analysis through projection onto first 2 PCs, Scree plot eigenvalue comparison, and sine of eigenvectors. (C) KNN/SNN analysis through SNN neighborhood Jaccard index and degree ratio (v7/v6) per cell. (D) Clustering and UMAP analysis through UMAP plots of each condition, with alluvial plot showing cluster assignment mapping and degree of agreement. (E) Differential expression analysis through overlap of all significant ($p < 0.05$) marker genes across all clusters.

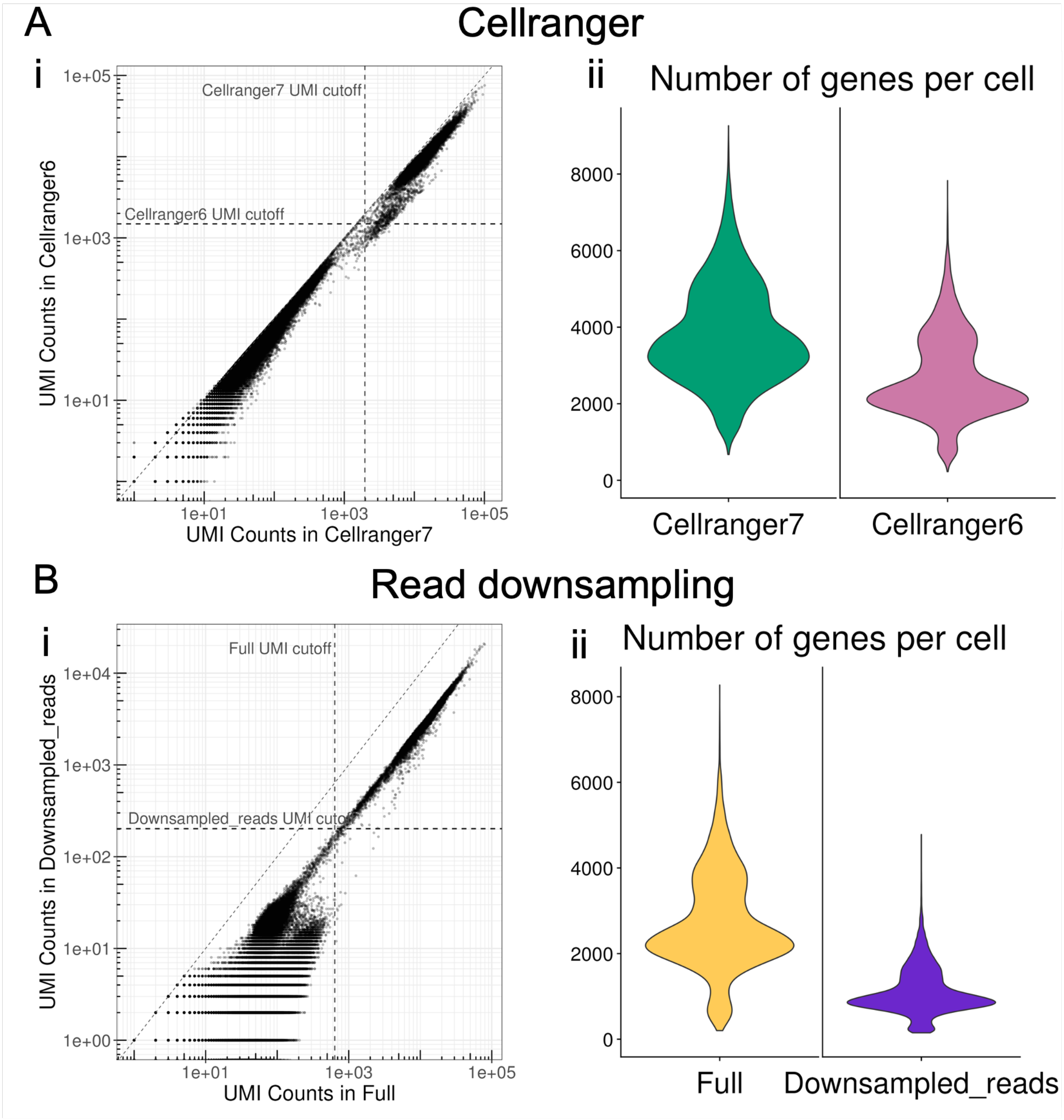


**Supplemental Figure 14:** UMI counts pre-filtering and number of features under different conditions. (A) Cell Ranger v7 vs. v6. (B) Full size dataset vs. downsampled reads to 4%.

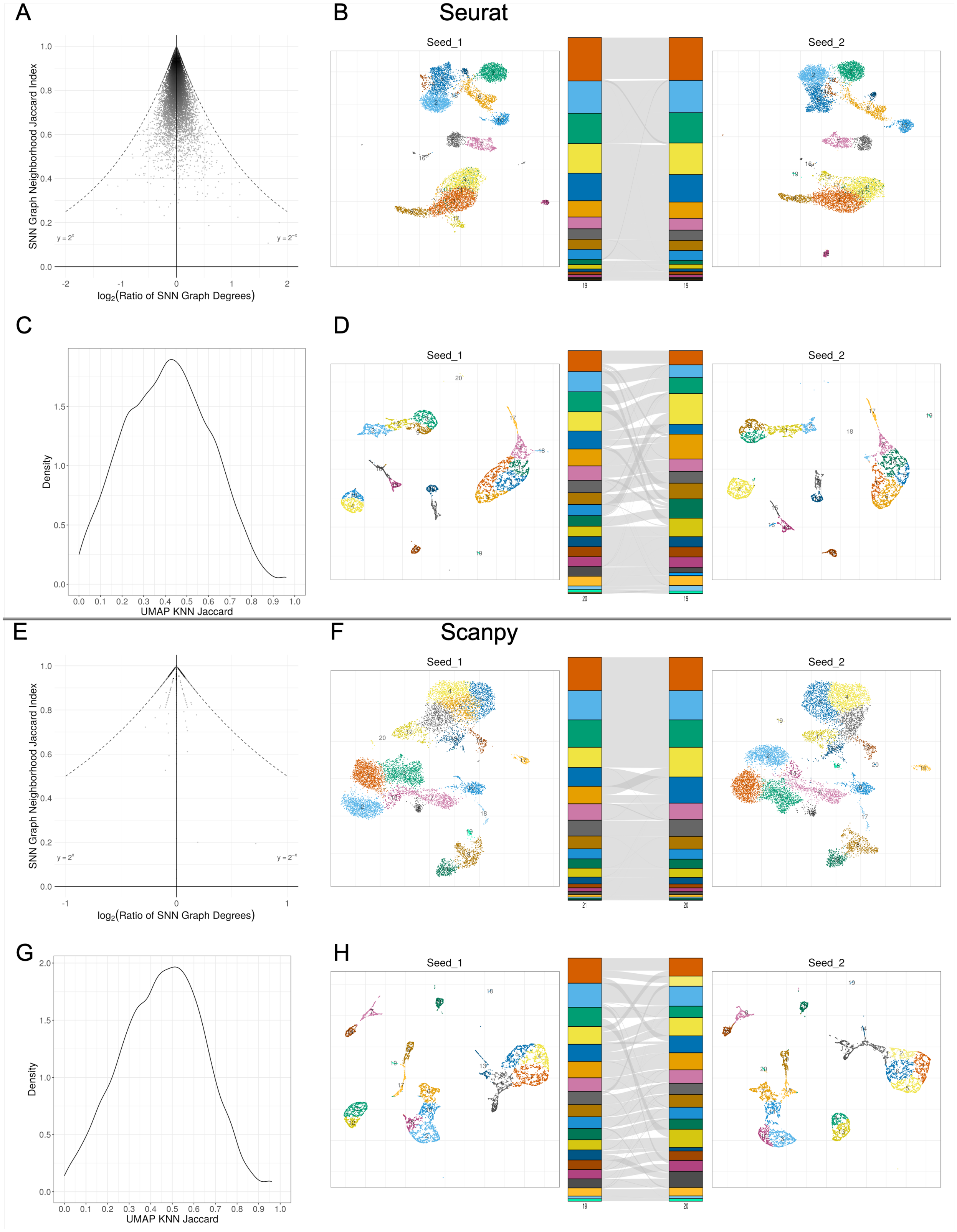
A
B
Seurat
C
D
E
F
Scanpy
G
H
Seed_1
Seed_2
SNN Graph Neighborhood Jaccard Index
log2(Ratio of SNN Graph Degrees)
Density
UMAP KNN Jaccard

**Supplemental Figure 15:** Variability introduced by differences in random seeds for functions involving randomness. (A) Seurat SNN graph generation (Annoy), with different seeds for KNN graph generation. (B) Seurat clustering (Louvain), with different seeds for Louvain; and UMAP-projection (uwot), with different seeds for UMAP projection. (C) Seurat KNN graph generation from UMAP space, with different seeds for UMAP projection. (D) Seurat clustering (Leiden) and UMAP projection from UMAP space, with different seeds for UMAP projection. (E) Scanpy SNN graph generation (NNDescent), with different seeds for KNN graph generation. (F) Scanpy clustering (Leiden), with different seeds for Leiden; and UMAP-projection (UMAP-learn), with different seeds for UMAP projection. (G) Scanpy KNN graph generation from UMAP space, with different seeds for UMAP projection. (H) Scanpy clustering (Leiden) and UMAP projection from UMAP space, with different seeds for UMAP projection.

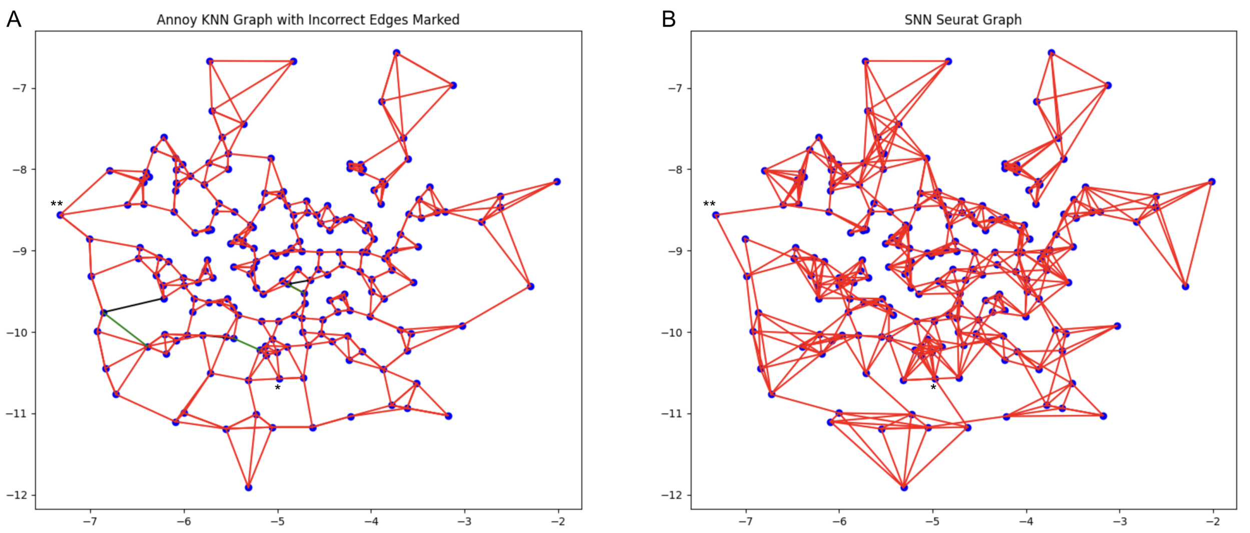


**Supplemental Figure 16:** Comparison of KNN and SNN graphs. (A) KNN graph (k=3) constructed with the approximate Annoy algorithm (directedness not shown). Black = incorrectly added edges; Green = incorrectly omitted edges. (B) SNN graph with Seurat's pruning method. An example hub node is marked with *. An example peripheral node is marked with **. Note: the SNN graph with Scanpy's method is simply the undirected KNN graph.

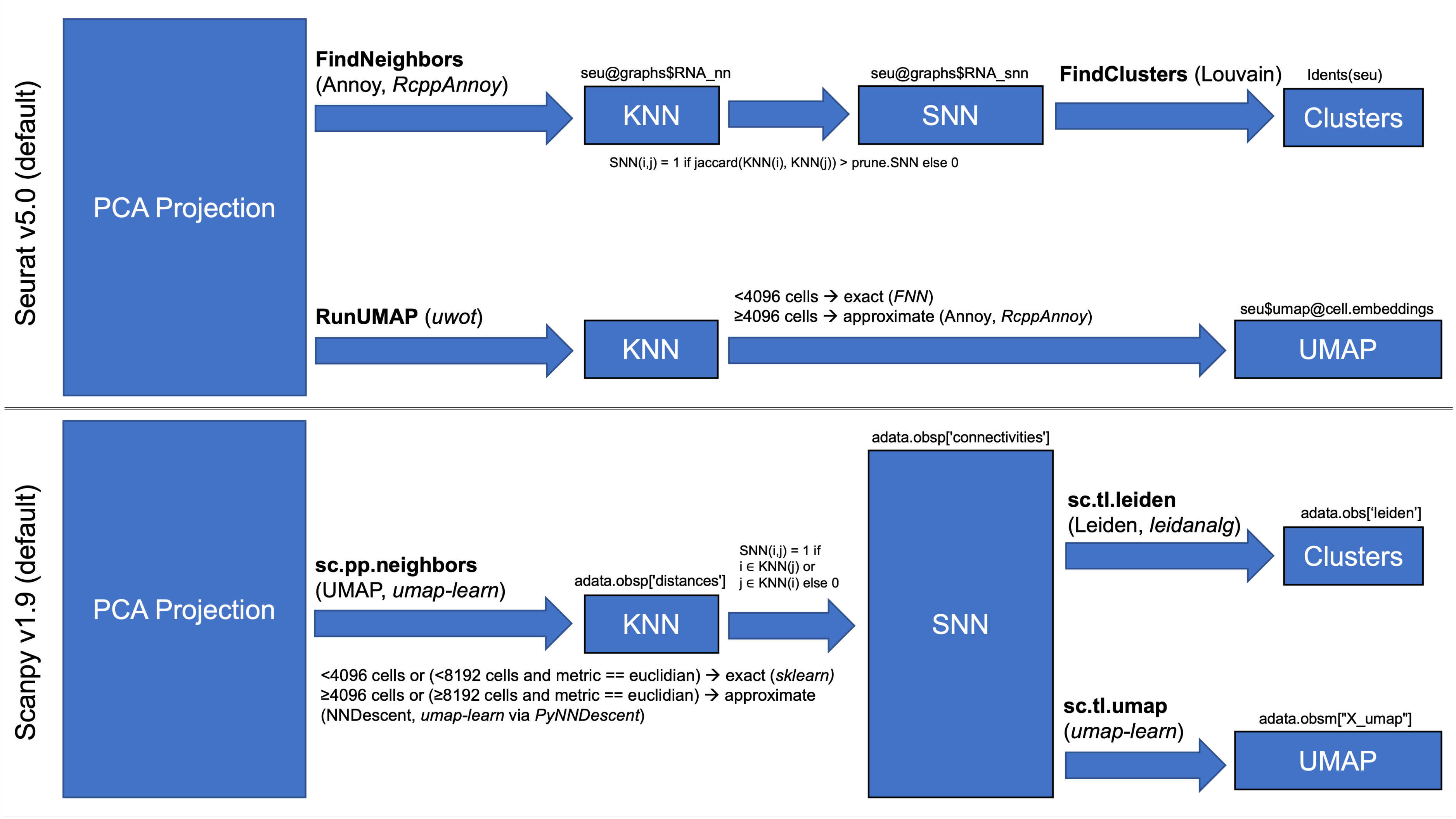
Seurat v5.0 (default)
PCA Projection
FindNeighbors
(Annoy, RcppAnnoy)
seu@graphs$RNA_nn
KNN
seu@graphs$RNA_snn
SNN
SNN(i,j) = 1 if jaccard(KNN(i), KNN(j)) > prune.SNN else 0
FindClusters (Louvain)
Idents(seu)
Clusters
RunUMAP (uwot)
KNN
<4096 cells → exact (FNN)
≥4096 cells → approximate (Annoy, RcppAnnoy)
seu$
UMAP
Scanpy v1.9 (default)
PCA Projection
sc.pp.neighbors
(UMAP, umap-learn)
adata.obsp['distances']
KNN
SNN(i,j) = 1 if
i ∊ KNN(j) or
j ∊ KNN(i) else 0
<4096 cells or (<8192 cells and metric == euclidian) → exact (sklearn)
≥4096 cells or (≥8192 cells and metric == euclidian) → approximate
(NNDescent, umap-learn via PyNNDescent)
adata.obsp['connectivities']
SNN
sc.tl.leiden
(Leiden, leidanalg)
adata.obs['leiden']
Clusters
sc.tl.umap
(umap-learn)
adata.obsm["X_umap"]
UMAP

**Supplemental Figure 17:** Schematic of protocol describing generation and downstream use of KNN and SNN graphs. Top is for Seurat, bottom is for Scanpy. KNN(i) refers to the full set of k-nearest neighbors for cell i. Bold = function/method name; Italics = outside package utilized.

| **Seurat v5.0.2** | **Filtering** | **Normalization** | **HVG selection** | **Regression, Scaling** | **PCA** | **SNN** | **Clustering** | **UMAP** | **DE** |
|---|---|---|---|---|---|---|---|---|---|
| **Function nam** | CreateSeuratObject | NormalizeData | FindVariableFeatures | ScaleData | RunPCA | FindNeighbors | FindClusters | RunUMAP | FindAllMarkers |
| **Defaults** | **min.cells = 0**<br>**min.features = 0** | *scale.factor = 10000*<br>*normalization.method = "LogNormalize"* | selection.method = "vst"<br>**loess.span = 0.3**<br>*clip.max = "auto"*<br>*mean.function = FastExpMean*<br>*dispersion.function = FastLogVMR*<br>**num.bin = 20**<br>*binning.method = "equal_width"*<br>nfeatures = 2000<br>mean.cutoff = c(0.1, 8)<br>dispersion.cutoff = c(1, Inf) | **vars.to.regress = NULL**<br>*latent.data = NULL*<br>*split.by = NULL*<br>*model.use = "linear"*<br>*use.umi = FALSE*<br>*do.scale = TRUE*<br>**do.center = TRUE**<br>scale.max = 10 | **npcs = 50**<br>*rev.pca = FALSE*<br>*weight.by.var = TRUE*<br>*approx = TRUE*<br>**features = NULL (HVGs)** | *reduction = "pca"*<br>dims = 1:10<br>k.param = 20<br>*prune.SNN = 1/15*<br>*nn.method = "annoy"*<br>*n.trees = 50*<br>**annoy.metric = "euclidean"**<br>nn.eps = 0<br>*l2.norm = FALSE* | *modularity.fxn = 1*<br>*initial.membership = NULL*<br>*node.sizes = NULL*<br>resolution = 0.8<br>method = "matrix"<br>algorithm = 1 (original Louvain)<br>*n.start = 10*<br>n.iter = 10<br>*group.singletons = TRUE* | umap.method = "uwot"<br>*n.neighbors = 30L*<br>**n.components = 2L**<br>*metric = "cosine"*<br>**n.epochs = NULL**<br>**learning.rate = 1**<br>min.dist = 0.3<br>**spread = 1**<br>*set.op.mix.ratio = 1*<br>*local.connectivity = 1L*<br>**repulsion.strength = 1**<br>**negative.sample.rate = 5**<br>a = NULL<br>b = NULL<br>dens.lambda = 2<br>dens.frac = 0.3<br>dens.var.shift = 0.1 | *logfc.threshold = 0.1*[a]<br>test.use = "wilcox"<br>*min.pct = 0.01*[b]<br>*min.diff.pct = -Inf*<br>*only.pos = FALSE*<br>*max.cells.per.ident = Inf*<br>*latent.vars = NULL*<br>*min.cells.feature = 3*<br>*min.cells.group = 3*<br>*mean.fxn = NULL*<br>*return.thresh = 0.01*<br>*base = 2* |
| **(non-default) Arguments to match Scanpy defaults** | | | selection.method = "mean.var.plot"<br>mean.cutoff = c(0.0125, 3)<br>dispersion.cutoff = c(0.5, Inf) | vars.to.regress = c("nCount_RNA", "pct_mt")<br>scale.max = Inf | | k.param = 15<br>dims = *total_pcs_used* | algorithm = "leiden"<br>resolution = 1.0 | umap.method = "umap-learn"<br>min.dist = 0.5 | logfc.threshold= 0<br>min.pct = 0<br>return.thresh = 1.0<br>min.cells.group = 1 |

[a]default changed from Seurat v4 (was logfc.threshold = 0.25)
[b]default changed from Seurat v4 (was min.pct = 0.1)

**Supplemental Table 1:** Seurat function details for scRNA-seq pipeline agreement with Scanpy. Bold = parameter in agreement by default; italics = no analogous paremeter in Scanpy; green = equivalent by default; yellow = equivalent with matched arguments; orange = partially incompatible, partially equivalent with matched arguments (depending on algorithm choice); red = incompatible

| **Scanpy v1.9.5** | **Filtering** | **Normalization** | **HVG selection** | **Regression, Scaling** | **PCA** | **SNN** | **Clustering** | **UMAP** | **DE** |
|---|---|---|---|---|---|---|---|---|---|
| **Function name** | sc.pp.filter_cells, sc.pp.filter_genes | sc.pp.normalize_total | sc.pp.highly_variable_genes | sc.pp.regress_out, sc.pp.scale | sc.tl.pca | sc.pp.neighbors | sc.tl.leiden, sc.tl.louvain | sc.tl.umap | sc.tl.rank_genes_groups |
| **Defaults** | **min_cells = None**<br>**min_genes = None**<br>*max_cells = None*<br>*max_genes = None*<br>*min_counts = None (genes)*<br>*min_counts = None (cells)*<br>*max_counts = None (genes)*<br>*max_counts = None (cells)* | *target_sum = None*<br>*exclude_highly_expressed = False*<br>*max_fraction = 0.05* | n_top_genes = None<br>min_mean = 0.0125<br>max_mean = 3<br>min_disp = 0.5<br>max_disp = inf<br>**span = 0.3**<br>**n_bins = 20**<br>flavor = "seurat" | **zero_center = True**<br>max_value = None | **n_comps = 50** or 1 – minimum dimension size<br>zero_center = True<br>*svd_solver = "arpack"*<br>**use_highly_variable = None (HVGs)** | n_neighbors = 15<br>n_pcs = None<br>*use_rep = None*<br>*knn = True*<br>*method = "umap"*<br>**metric = "euclidian"** | resolution = 1<br>*restrict_to = None*<br>*directed = True*<br>*use_weights = True*<br>n_iterations = -1<br>*partition_type = None*<br>*flavor = "vtraag" (louvain)* | min_dist = 0.5<br>**spread = 1.0**<br>*n_components = 2*<br>**maxiter = None**<br>**alpha = 1.0**<br>**gamma = 1**<br>**negative_sample_rate = 5**<br>*init_pos = "spectral"*<br>**a = None**<br>**b = None**<br>method = "umap" | *use_raw = None*<br>*reference = "rest"*<br>n_genes = None<br>rankby_abs = False<br>pts = False<br>method = None<br>corr_method = 'benjamini-hochberg'<br>*tie_correct = False* |
| **(non-default) Arguments to match Seurat defaults** | | | n_top_genes = 2000<br>flavor = "seurat_v3" | Omit sc.pp.regress_out<br>max_value = 10 | zero_center = False | n_neighbors = 20<br>n_pcs = 10<br>use_rep = 'X_pca' | function sc.tl.louvain instead of sc.tl.leiden<br>resolution = 0.8<br>n_iterations = 10 | min_dist = 0.3 | use_raw=True<br>pts = True<br>method = "wilcoxon"<br>corr_method = "bonferroni"<br>tie_correct = True<br>manual filtering (abs(log_fc) ≥ 0.1, pts ≥ 0.01, pts_rest ≥ 0.01 p_value < 0.01) |

**Supplemental Table 2:** Scanpy function details for scRNA-seq pipeline agreement with Seurat. Bold = parameter in agreement by default; italics = no analogous paremeter in Seurat; green = equivalent by default; yellow = equivalent with matched arguments; orange = partially incompatible, partially equivalent with matched arguments (depending on algorithm choice); red = incompatible
